# Commensal lifestyle regulated by a negative feedback loop between *Arabidopsis* ROS and the bacterial T2SS

**DOI:** 10.1101/2023.05.09.539802

**Authors:** Frederickson Entila, Xiaowei Han, Akira Mine, Paul Schulze-Lefert, Kenichi Tsuda

**Affiliations:** National Key Laboratory of Agricultural Microbiology, Hubei Hongshan Laboratory, Hubei Key Lab of Plant Pathology, College of Plant Science and Technology, Huazhong Agricultural University, Wuhan 430070, China; Department of Plant Microbe Interactions, Max Planck Institute for Plant Breeding Research, Carl-von-Linne-Weg 10, Cologne 50829, Germany; Shenzhen Institute of Nutrition and Health, Huazhong Agricultural University, Wuhan 430070, China.; Shenzhen Branch, Guangdong Laboratory of Lingnan Modern Agriculture, Genome Analysis Laboratory of the Ministry of Agriculture and Rural Affairs, Agricultural Genomics Institute at Shenzhen, Chinese Academy of Agricultural Sciences, Shenzhen, Guangdong 518120, China; JST PRESTO, Kawaguchi-shi, Saitama 332-0012, Japan; Laboratory of Plant Pathology, Graduate School of Agriculture, Kyoto University, Kyoto 606-8502, Japan

## Abstract

Despite the plant health-promoting effects of plant microbiota, these assemblages also comprise potentially detrimental microbes. How plant immunity controls its microbiota to promote plant health under these conditions remains largely unknown. We found that commensal bacteria isolated from healthy *Arabidopsis* plants trigger diverse patterns of reactive oxygen species (ROS) production via the NADPH oxidase RBOHD that selectively inhibited specific commensals, notably *Xanthomonas* L148. Through random mutagenesis, we found that L148 *gspE*, encoding a type II secretion system (T2SS) component, is required for the damaging effects of *Xanthomonas* L148 on *rbohD* mutant plants. *In planta* bacterial transcriptomics revealed that RBOHD suppresses most T2SS gene expression including *gspE*. L148 colonization protected plants against a bacterial pathogen, when *gspE* was inhibited by ROS or mutation. Thus, a negative feedback loop between *Arabidopsis* ROS and the bacterial T2SS tames a potentially detrimental leaf commensal and turns it into a microbe beneficial to the host.

## Introduction

In nature, plants host diverse microbes called the plant microbiota^1^. While the plant microbiota collectively contributes to plant health, they comprise microorganisms ranging from mutualistic to commensal, and pathogenic microbes^2^. The property of microbes as mutualistic, commensal, and pathogenic depends on the host and environmental condition^3–4^. Thus, the plant microbiota is not simply a collection of beneficial microbes, but various factors affect the property of microbes within the plant microbiota, which consequently determines plant health.

Upon recognition of microbial molecules, plants activate a battery of immune responses^5^. In the first layer of immunity, known as pattern-triggered immunity (PTI), plasma membrane-localized pattern recognition receptors (PRRs) recognize microbe-associated molecular patterns (MAMPs). For instance, the PRR FLAGELLIN SENSING 2 (FLS2) and EF-TU RECEPTOR (EFR) sense the bacteria-derived oligopeptides flg22 and elf18, respectively, in *Arabidopsis thaliana*^6, 7^. BRI1-ASSOCIATED RECEPTOR KINASE 1 (BAK1) and its close homolog BAK1-LIKE 1 (BKK1) function as co-receptors for LRR-RLK-type PRRs such as FLS2 and EFR^8^. The LysM-RLK CHITIN ELICITOR RECEPTOR KINASE 1 (CERK1) is an essential co-receptor for fungal chitin and bacterial peptidoglycans^9^. Activated PRRs trigger various immune responses such as the production of reactive oxygen species (ROS), calcium influx, MAP kinase activation, transcriptional reprogramming, and the production of defense phytohormones and specialized metabolites^10^. PTI contributes not only to pathogen resistance but also to the maintenance of healthy microbiota as evidenced by dysbiosis and disease symptoms observed on leaves of *A. thaliana* genotypes with severely impaired PTI responses^11, 12^. However, the molecular mechanism by which PTI-associated immune responses regulate microbial pathogens and maintain healthy microbiota remains unclear.

One prominent PTI output involves activation of the plasma membrane-localized NADPH oxidase RESPIRATORY BURST OXIDASE HOMOLOG D (RBOHD), which produces the ROS O_2-_ in the extracellular space, which can then be readily converted to H_2_O_2_ via superoxide dismutase in the apoplast^13^. Extracellular ROS can be sensed by a plasma membrane-localized sensor and can be translocated into the cell to mediate plant immune responses^14^. Extracellular ROS can also directly exert cellular toxicity on microbes^15^. ROS functions in regulating not only resistance against pathogens, but also the composition and functions of the plant microbiota. For instance, RBOHD-mediated ROS production inhibits Pseudomonads in the *A. thaliana* rhizosphere^16^. ROS also prohibits dysbiosis in *A. thaliana* leaves by suppressing *Xanthomonas*^17^. Plant RBOHD-mediated ROS induces the production of the phytohormone auxin in the beneficial bacterium *Bacillus velezensis* and promotes root colonization in *A. thaliana*^18^. These studies exemplify the importance of RBOHD-mediated ROS production in the regulation of plant microbiota. However, how ROS specifically regulates microbial metabolism and growth remains unknown. Furthermore, while ROS exhibits general cell toxicity to organisms, not all microbes are sensitive to plant-produced ROS. For instance, the growth of the bacterial pathogen *Pseudomonas syringae* pv. *tomato* DC3000 (*Pto*) was not affected by mutation in *RBOHD* in *A. thaliana*^19^. This indicates that ROS exerts differential actions on microbes, but the basis for this selectivity needs to be explored.

Secretion systems are crucial for bacterial pathogens to efficiently infect the host plant through the secretion of effector proteins, among which the type III secretion system (T3SS) has been well documented as the essential pathogenicity component of many phytopathogenic bacteria^20^. The key function of T3SS is to introduce type III effectors (T3Es) directly into the host cell, thereby suppressing plant immunity and promoting virulence^20^. Some nitrogen-fixing rhizobacteria also utilize the T3SS to promote symbiosis with their legume host^21–22^. A number of T3Es have been identified to be recognized by the intracellular nucleotide-binding domain leucine-rich repeat receptors (NLRs), activating effector-triggered immunity^23^. These indicate the paramount significance of the T3SS for the interaction between host and bacteria. In addition to the T3SS, the type II secretion system (T2SS) has been shown to be necessary for the pathogenesis of many phytopathogenic bacteria and mainly functions to secrete enzymes to degrade host barriers and promote virulence^20^. Interestingly, the root commensal *Dyella japonica* MF79 requires the T2SS components *gspD* and *gspE* to release immune-suppressive factors that help the root colonization of a non-immune suppressive commensal in *A. thaliana*^24^. However, whether and how plant immunity controls T2SS activity of its microbiota remains unknown.

In this study, we investigated the impact of *A. thaliana* immune responses to commensal bacteria isolated from healthy *A. thaliana* plants with a focus on RBOHD-mediated ROS. Using a bacterial random mutagenesis screen and *in planta* bacterial transcriptomics, we revealed that RBOHD-mediated ROS directly suppresses the T2SS of a potentially harmful *Xanthomonas* L148, thereby converting *Xanthomonas* L148 into a commensal. Moreover, this “tamed” *Xanthomonas* increased host resistance against the bacterial pathogen *Pto*.

## Results

### Different commensal bacteria trigger diverse ROS production patterns via distinct mechanisms

We investigated variations in immune responses triggered by the colonization of different commensal bacteria in *A. thaliana* leaves with ROS production as the readout. First, we measured ROS production in leaves of *fls2*, *efr*, *cerk1*, *fls2 efr cerk1* (*fec*), *bak1 bkk1 cerk1* (*bbc*), and *rbohD* mutants as well as Col-0 wild-type plants in response to the MAMPs flg22, elf18, and chitin heptamer. ROS production was dependent on the corresponding (co)receptor and *RBOHD*, indicating the suitability of our experimental system (Figure 1a and Supplementary Figure S1). Next, we measured ROS production in leaves of the same mutant panel in response to taxonomically diverse 20 live and heat-killed commensal bacterial strains that were previously isolated from healthy *A. thaliana* leaves and roots as well as soil^25^ and that were used for plant-bacterial co-transcriptomics^26^ (Figure 1d). These commensal bacteria triggered diverse ROS production patterns. For instance, both live and heat-killed *Pseudomonas* L127 triggered ROS production with the heat-killed bacteria eliciting stronger ROS production, which is a general trend for all commensal bacterial strains (Figure 1c and Supplementary Figure S2). On the other hand, only heat-killed but not live *Burkholderia* L177 triggered ROS production, suggesting that L177 possesses MAMP(s) that are potentially recognized by plants but live L177 does not expose such MAMPs. Further, neither the live nor the heat-killed *Flavobacterium* R935 triggered ROS production. We observed neither obvious phylogenetic signatures predictive for the capability to induce ROS, nor of the tissue of origin from which these commensals were isolated. We also observed different dependencies of commensal bacteria-induced ROS on the MAMP (co)receptors. For instance, ROS production by both live and heat-killed *Exiguobacterium* L187 was dependent on *EFR* but not *FLS2* and *CERK1*. This *EFR* dependency for commensal bacteria-induced ROS production was observed for other strains, but we detected no or only weak effects of mutations in *FLS2* and *CERK1*. These results suggest that the recognition of EF-Tu-derived peptides via EFR is the primary mechanism for ROS production by commensal bacteria in *A. thaliana* leaves. However, there were commensal bacteria such as *Pseudomonas* L127 that stimulated ROS in *fec* and *bbc* mutant plants (Supplementary Figure S2), indicating that MAMPs other than flg22, elf18, and peptidoglycans are responsible for ROS production induced by commensal bacteria in some cases.

**Figure 1.**
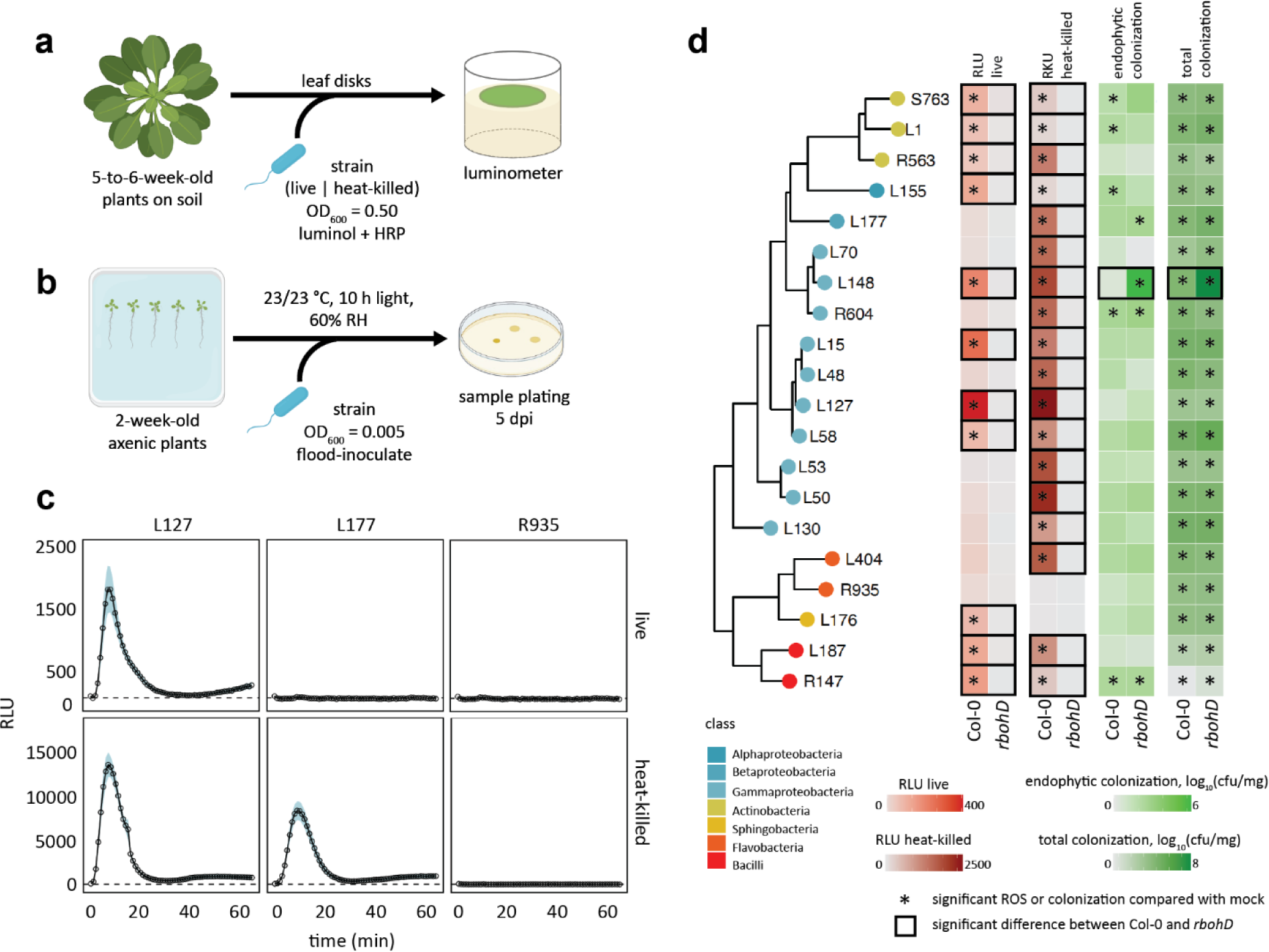
Immunogenic and colonization profile of microbiota members in mono-associations. Schematic diagram of ROS burst assay (**a**) in leaf discs from 5 to 6-week-old Col-0 plants treated with live or heat-killed bacterial cells (OD600=0.5) and colonization capacities (**b**) of the microbiota members upon flood inoculation (OD600=0.005) of 2-week-old Col-0 plants at 5 dpi. **c**, ROS burst profile of representative strains with varying behaviors: immune-active,-evasive, and -quiescent, for *Pseudomonas* L127, *Burkholderia* L177, and *Flavobacterium* R935, respectively (see Supplementary Figure S2 for the full ROS burst profiles). **d**, Phylogenetic relationship of the selected microbiota members and the heatmap representation of their corresponding ROS burst profiles using live and heat-killed cells, and their respective colonization capacities in leaves of Col-0 and *rbohD* plants; * indicates significant within-genotype difference of the trait between mock and the bacterial strain in question; □ indicates significant within-strain difference of the trait between Col-0 and *rbohD* plants (ANOVA with *post hoc* Tukey’s test, *P* ≤ 0.05). Experiments were repeated at least two times each with 8 biological replicates for ROS assay and 3–4 biological replicates for colonization assays (See Supplementary Figure S3 for the full colonization profiles and Supplementary Table S2 for detailed descriptions of the strains included). Some illustrations were created with BioRender.

### Plant-derived ROS differentially affects the colonization of commensal bacteria

We found that ROS production by all live and heat-killed commensal bacteria was completely dependent on *RBOHD*, indicating that RBOHD is mainly responsible for plant ROS production triggered by these commensal bacteria. Plant-produced ROS via RBOHD can affect the colonization of commensal bacteria. We then determined total and endophytic bacterial titers of different commensals in leaves of Col-0 wild-type and *rbohD* as well as *fls2*, *efr*, *fec*, and *bbc* mutant plants. We grew plants on agar plates for 14 days and flood-inoculated with individual commensal bacteria followed by the determination of bacterial titer (Figure 1b, Supplementary Figure S3). To our surprise, while we did observe increased colonization of some commensal bacteria in some of the MAMP (co)receptor mutants compared with Col-0 wild-type plants, we were largely unable to detect any impact of the *rbohD* mutation on either total or endophytic commensal colonization (Figure 1d and Supplementary Figure S3). Also, there is no significant relationship between the ROS immunogenicity and the colonization capacity of the commensal bacteria (Supplementary Figure S4). These findings suggest that *Arabidopsis* recognizes commensal bacteria and produces ROS that does not have a detectable impact on most commensal bacterial colonization, at least in mono-associations. By contrast, both total and endophytic colonization of *Xanthomonas* L148 was dramatically increased in *rbohD* mutant compared with Col-0 wild-type plants (Figure 1d and Supplementary Figure S3), suggesting that RBOHD-mediated ROS suppresses *Xanthomonas* L148 colonization, consistent with a recent finding^17^.

### *Xanthomonas* L148 is detrimental to *rbohD* mutant but not Col-0 wild-type plants

Leaf colonization of *rbohD* mutant plants with live *Xanthomonas* L148 led to host mortality within 5 days post inoculation (dpi), in contrast to asymptomatic wild-type Col-0 plants (Figure 2a). In an orthogonal system, we infiltrated leaves with *Xanthomonas* L148 and observed disease-like symptoms only in *rbohD* after 3 dpi (Figure 2b-d). As *Xanthomonas* L148 activated ROS burst in Col-0 leaves, but not in *rbohD*, *Xanthomonas* L148 pathogenicity might be suppressed by the ROS pathway (Figure 2e). Furthermore, *Xanthomonas* L148 not only persisted on the leaf surface but aggressively colonized the apoplast of *rbohD* mutants compared with Col-0 at 3 dpi (Figure 2f). Together, *Xanthomonas* L148 is potentially pathogenic and its deleterious effect depends on the absence of *RBOHD*.

**Figure 2.**
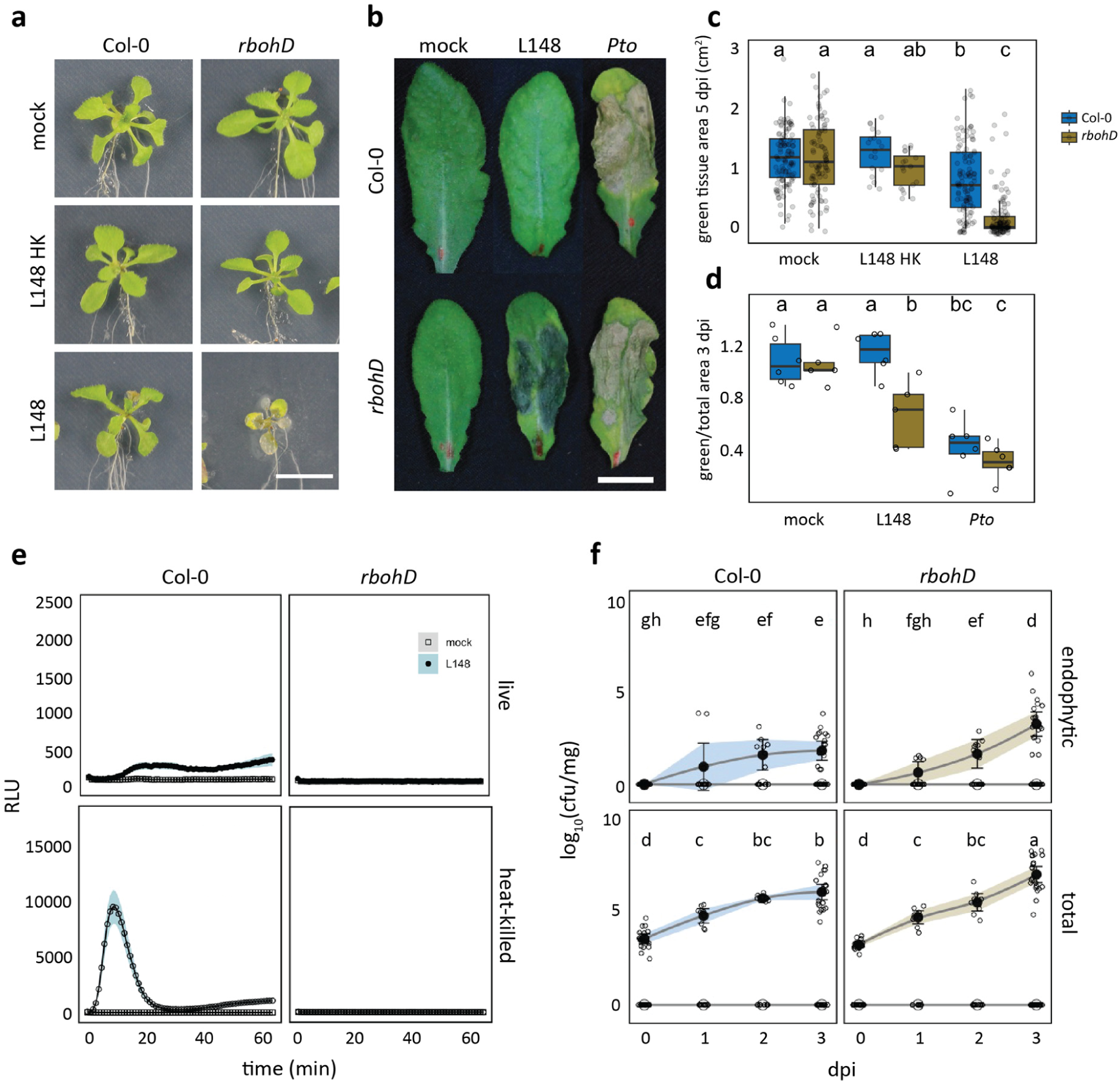
*Xanthomonas* L148 is detrimental to *rbohD* mutant but not to Col-0 wild-type plants. **a**, **c**. Representative images (**a**) and quantification of green tissue area (**c**) as the plant health parameter. 14-day-old Col-0 and *rbohD* plants grown on agar plates were flood-inoculated with mock and live and heat-killed (HK) *Xanthomonas* L148 (OD600=0.005). Samples were taken at 5 dpi (4 independent experiments each with at least 5 biological replicates). **b, d.** Representative images **(b)** and quantification of percentage green tissue of leaves **(d)** hand-infiltrated with mock, *Xanthomonas* L148 and *Pto* (OD600=0.2). Samples were taken at 3 dpi (2 independent experiments each with 3–4 biological replicates). **e**, ROS burst profile of leaf discs of 5–6-week-old Col-0 and *rbohD* plants treated with live and heat-killed *Xanthomonas* L148 (OD600=0.5) (at least 4 independent experiments each with 8 biological replicates). **f**, Infection dynamics of *Xanthomonas* L148 upon flood inoculation of 14-day-old Col-0 and *rbohD* plants grown in agar plates (OD600=0.005). Leaf samples were harvested at 0 to 3 dpi for total and endophytic compartments (2 independent experiments each with 3–4 biological replicates). Results in **c** and **d** are depicted as box plots with the boxes spanning the interquartile range (IQR, 25^th^ to 75^th^ percentiles), the mid-line indicates the median, and the whiskers cover the minimum and maximum values not extending beyond 1.5x of the IQR. Results in **f** are shown as line graphs using Locally Estimated Scatter Plot Smoothing (LOESS) with error bars and shadows indicating the standard errors of the mean. **c,d,f**, ANOVA with *post hoc* Tukey’s test. Different letters indicate statistically significant differences (*P* ≤ 0.05).

### *Xanthomonas* L148 is largely insensitive to ROS *in vitro*

Due to their highly reactive nature, ROS can oxidize bacterial components, which can lead to extensive cellular damage. This might explain why *Xanthomonas* L148 is pathogenic to *rbohD* mutant but not to Col-0 wild-type plants. We tested the sensitivity of *Xanthomonas* L148 to ROS compounds by instantaneous *in vitro* exposure to H_2_O_2_ or O_2-1_. To our surprise, *Xanthomonas* L148 seemed to tolerate acute treatments with ROS and retained viability up to ROS concentrations of 1 mM (Supplementary Figure S5a-b). Similar findings were obtained when a ROS-generating compound, paraquat (PQ, Supplementary Figure S5c), was used. It can be argued that the adverse effects of ROS *in vitro* can only be observed upon continuous ROS treatment. However, we did not observe any significant effects on the growth rates of *Xanthomonas* L148 upon chronic exposure to PQ (Supplementary Figure S5d). This suggests that the rampant proliferation of *Xanthomonas* L148 in *rbohD* plants is not due to the direct microbiocidal effects of ROS but other mechanisms.

### *Xanthomonas* L148 pathogenic potential is partially suppressed by the presence of other leaf microbiota members

*Xanthomonas* L148 was isolated from macroscopically healthy *A. thaliana* plants grown in their natural habitat, indicating that it is a constituent of the native leaf microbiota of *A*. *thaliana*. While *Xanthomonas* L148 was detrimental to *rbohD* mutant plants in a mono-association condition, it can be postulated that in a microbial community setting, *Xanthomonas* L148 is disarmed and *rbohD* plants become asymptomatic. To test this, we constructed a synthetic bacterial community which consists of nine leaf-derived isolates that were found to be robust leaf colonizers and cover the major phyla of the native bacterial microbiota of leaves^27–29^, which we refer to as LeafSC (Supplementary Figure S6b, please see Supplementary Table S2 for the strain details). We also assessed the dose-dependency of the disease onset by using different proportions of *Xanthomonas* L148 in relation to the entire LeafSC, with L148_P1_ as a dose equivalent to that of each synthetic community member (*Xanthomonas* L148/LeafSC, 1:9), while L148_P9_ is a dosage that is equal to the entire bacterial load of the synthetic community (*Xanthomonas* L148/LeafSC, 9:9). Flood inoculation of plants with the LeafSC did not result in any observable disease symptoms (Supplementary Figure S6a and S6c). As expected, inoculation with *Xanthomonas* L148 resulted in substantial mortality of *rbohD* plants compared with Col-0 wild-type plants. The killing activity of *Xanthomonas* L148 was somewhat reduced in *rbohD* plants when other microbiota strains were present, but this counter effect was overcome when a higher dose of *Xanthomonas* L148 was used (Supplementary Figure S6a and S6c). These findings imply that a functional leaf microbiota contributes to the partial mitigation of disease symptoms caused by *Xanthomonas* L148 in *rbohD* plants, possibly through niche occupancy, resource competition, or antibiosis.

### *Xanthomonas* L148::Tn5 mutant screening unveils genetic determinants of its pathogenic potential

*Xanthomonas* L148 is a conditional pathogen and its virulence is unlocked in the absence of *RBOHD* in the plant host. We aimed to identify the bacterial genetic determinants of this trait through a genome-wide mutant screening. We developed and optimized a robust high-throughput screening protocol (Figure 3a, Supplementary Figure S7a) and generated and validated a *Xanthomonas* L148 Tn5 mutant library (Supplementary Figure S7b–d). Using the high-throughput protocol, this Tn5 mutant library was phenotyped for the loss-of-*rbohD* mortality. From 6,862 transposon insertional mutants, 214 candidate strains consistently failed to exert pathogenicity on *rbohD* mutant plants (Figure 3b, See Supplementary Dataset S1 for the complete list of the candidate mutant strains). Most of the 214 strains did not exhibit significant defects in their *in vitro* growth parameters (growth rate, biofilm formation, and motility) in rich TSB medium or minimal XVM2 medium (Figure 3c). We found that out of the 214 strains, only 124 had transposon insertions in genes with functional annotations. These strains were retested in a square plate agar format, and 18 bacterial mutants exhibited consistent loss-of-*rbohD* mortality. Out of these 18 strains, three showed very strong phenotypes, namely *gspE*::Tn5, *alaA*::Tn5, and *rpfF*::Tn5 (Figure 3d-f). The candidate gene *gspE* encodes a core ATPase component of the T2SS; *alaA* encodes an alanine-synthesizing transaminase involved in amino acid metabolism; and *rpfF* encodes a synthase for diffusible signaling factor (DSF), a constituent of the quorum sensing machinery in bacteria (Figure 3d).

**Figure 3.**
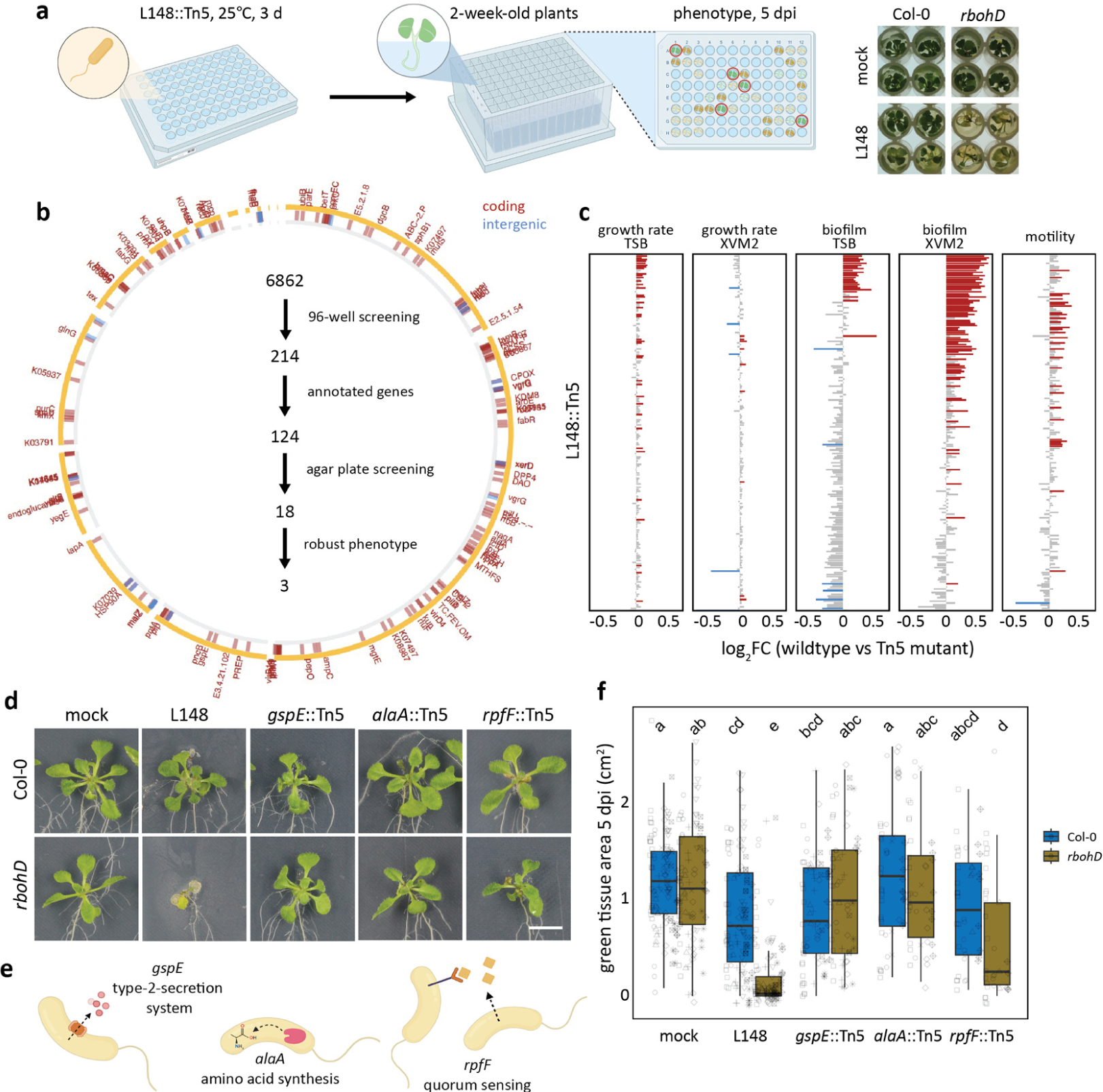
*Xanthomonas* L148::Tn5 mutant screening unveils genetic determinants of its pathogenic potential. **a**, Schematic diagram of the optimized high-throughput genetic screening for the *Xanthomonas* L148::Tn5 mutant library. Bacterial strains were inoculated onto 2-week-old *rbohD* plants followed by phenotyping at 5 dpi. **b**, Genomic coordinates of genes disrupted in the 214 *Xanthomonas* L148::Tn5 candidate strains. A total of 6,862 *Xanthomonas* L148::Tn5 strains were screened for loss of *rbohD* killing activity in a 96-well high-throughput format (2 independent experiments). We identified 124 strains with functional annotations, which were subsequently screened using the agar plate format, resulting in 18 strains with robust phenotypes. Finally, 3 strains were selected as the best-performing candidate strains. **c**, *In vitro* phenotypes of the 214 candidate strains: growth rates, biofilm production, and motility in rich TSB medium; growth rates and biofilm production in a minimal XVM2 medium. Data from 2 independent experiments each with 2–3 biological replicates were used for ANOVA with a *post hoc* Least Significant Difference (LSD) test. Red and blue bars indicate significantly higher or lower than the wild-type *Xanthomonas* L148 (*P* ≤ 0.05), respectively. **d**, **f**. Representative images **(d)** and quantification of green tissue area **(f)** as plant health parameter of Col-0 and *rbohD* plants flood mono-inoculated with *Xanthomonas* L148::Tn5 strains (OD600=0.005). Samples were harvested at 5 dpi. Data from at least 4 independent experiments each with 3–4 biological replicates were used for ANOVA with a *post hoc* Tukey’s test. Different letters indicate statistically significant differences (*P* ≤ 0.05). **e**, Graphical representation of the functions of the candidate genes. Results in **f** are depicted as box plots with the boxes spanning the interquartile range (IQR, 25^th^ to 75^th^ percentiles), the mid-line indicates the median, and the whiskers cover the minimum and maximum values not extending beyond 1.5x of the IQR. Some of the illustrations were created using BioRender.

### T2SS, amino acid metabolism, and quorum sensing underpin the conditional pathogenicity of *Xanthomonas* L148

We re-evaluated the candidate mutant strains using leaf-infiltration assays. The results showed that the disease progression required live *Xanthomonas* L148 as heat-killed bacteria did not elicit the same response (Figure 4a). Consistent with the previous systems (high-throughput and square plate set-ups), the mutant strains lost their capacity to cause disease symptoms on *rbohD* mutant plants (Figure 4a). As shown before, wild-type *Xanthomonas* L148 exhibited increased colonization in both total and endophytic compartments of *rbohD* leaves. By contrast, *gspE*::Tn5 mutant exhibited colonization capacities comparable to *Xanthomonas* L148 wild-type in Col-0 leaves, but failed to colonize to the same level on *rbohD* plants (Figure 4b). On the other hand, *alaA*::Tn5 mutants had a compromised colonization capacity in Col-0 plants, while *rpfF*::Tn5 mutant strains colonized *rbohD* leaves to a similar extent to wild-type *Xanthomonas* L148. Nonetheless, all of the mutant strains not only persisted but were able to actively colonize the leaf endosphere (Figure 4b). This indicates that *gspE*::Tn5 mutant retains its overall colonization ability, while its capacity to efficiently colonize *rbohD* plants is specifically compromised compared to wild-type L148. Correlation analysis revealed a negative relationship between host colonization and plant health, indicating that the observed leaf symptoms can be explained by the aggressive colonization of the wild-type strain (Figure 4d).

**Figure 4.**
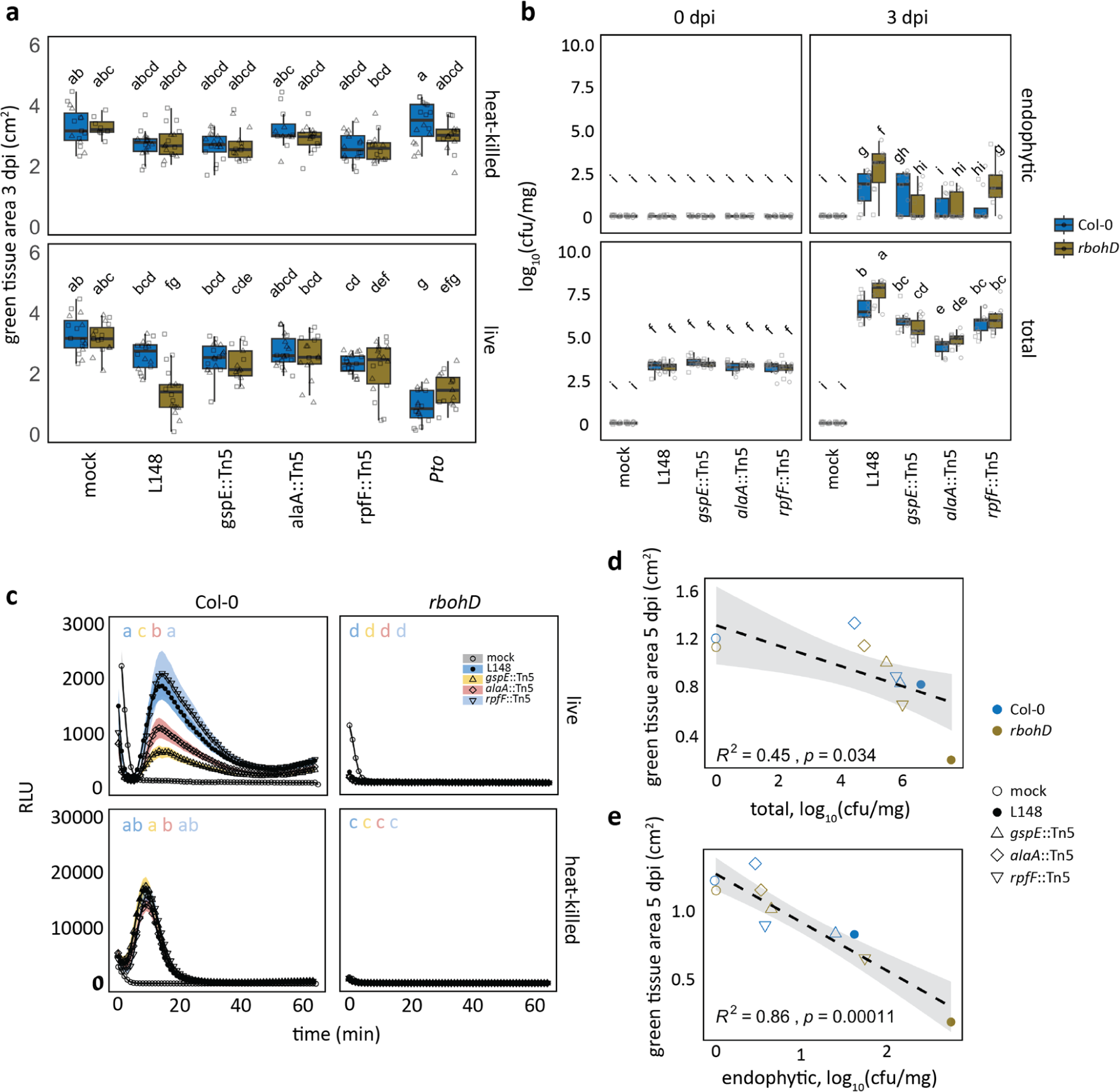
T2SS, amino acid metabolism, and quorum sensing underpin conditional pathogenicity of *Xanthomonas* L148 in *rbohD* plants. **a**, Quantification of green tissue area of hand-infiltrated 5 to 6-week-old Col-0 and *rbohD* leaves with *Xanthomonas* L148::Tn5 mutant strains using live and heat-killed cells as inoculum (OD600=0.2). Samples were collected at 3 dpi (2 independent experiments each with 3–4 biological replicates). **b**, Infection dynamics in axenic Col-0 and *rbohD* plants flood-inoculated with *Xanthomonas* L148::Tn5 mutant strains (OD600=0.005). Samples were harvested at 0 to 3 dpi for total and endophytic leaf compartments (2 independent experiments each with 3–4 biological replicates). **a,b**, ANOVA with *post hoc* Tukey’s test. Different letters indicate statistically significant differences (*P* ≤ 0.05). Results in **a** and **b** are depicted as box plots with the boxes spanning the interquartile range (IQR, 25^th^ to 75^th^ percentiles), the mid-line indicates the median, and the whiskers cover the minimum and maximum values not extending beyond 1.5x of the IQR. **c**, ROS burst profile of leaf discs of 5–6-week-old Col-0 and *rbohD* plants treated with live and heat-killed *Xanthomonas* L148 wild-type and L148::Tn5 mutant strains (OD600=0.5) (at least 4 independent experiments each with 8 biological replicates). **d,e**, Pearson correlation analyses of plant health performance measured as green tissue area against bacterial colonization capacities in the total **(d)** and endophytic **(e)** compartments (R^2^, coefficient of determination).

None of the three mutant strains were defective in growth, biofilm production, or motility in rich TSB medium (Figure 5a-c). Also, the mutant strains remained insensitive to PQ treatment, indicating retained tolerance to chronic ROS exposure (Figure 5a). *In vitro* growth phenotypes were also unchanged in minimal XVM2 medium apart from an increase in biofilm production for *gspE*::Tn5 and *alaA*::Tn5 mutant strains (Figure 5d). Secretion of extracellular enzymes acting on plant cell walls is a canonical strategy used by plant pathogens to breach the host’s physical barriers^20^. Bacterial pathogens often utilize T2SS to deliver these enzymes into the apoplast of their plant host^30^. We conducted enzyme secretion plate assays to test the proficiency of these strains to degrade different substrates (carbohydrates, protein, and lipids). Wild-type *Xanthomonas* L148 was able to secrete extracellular enzymes that can degrade the proteinaceous compounds gelatin and non-fat dry milk and the carbohydrates pectin and carboxymethyl-cellulose. Notably, *gspE*::Tn5 mutant could not degrade these substrates in contrast to the wild-type and the other mutant strains, indicating impaired secretion activities (Figure 5e-f). This suggests that the lack of disease progression in *rbohD* plants with the *gspE*::Tn5 mutant strain can be explained by its inability to secrete extracellular enzymes to degrade the host plant cell walls via the T2SS.

**Figure 5.**
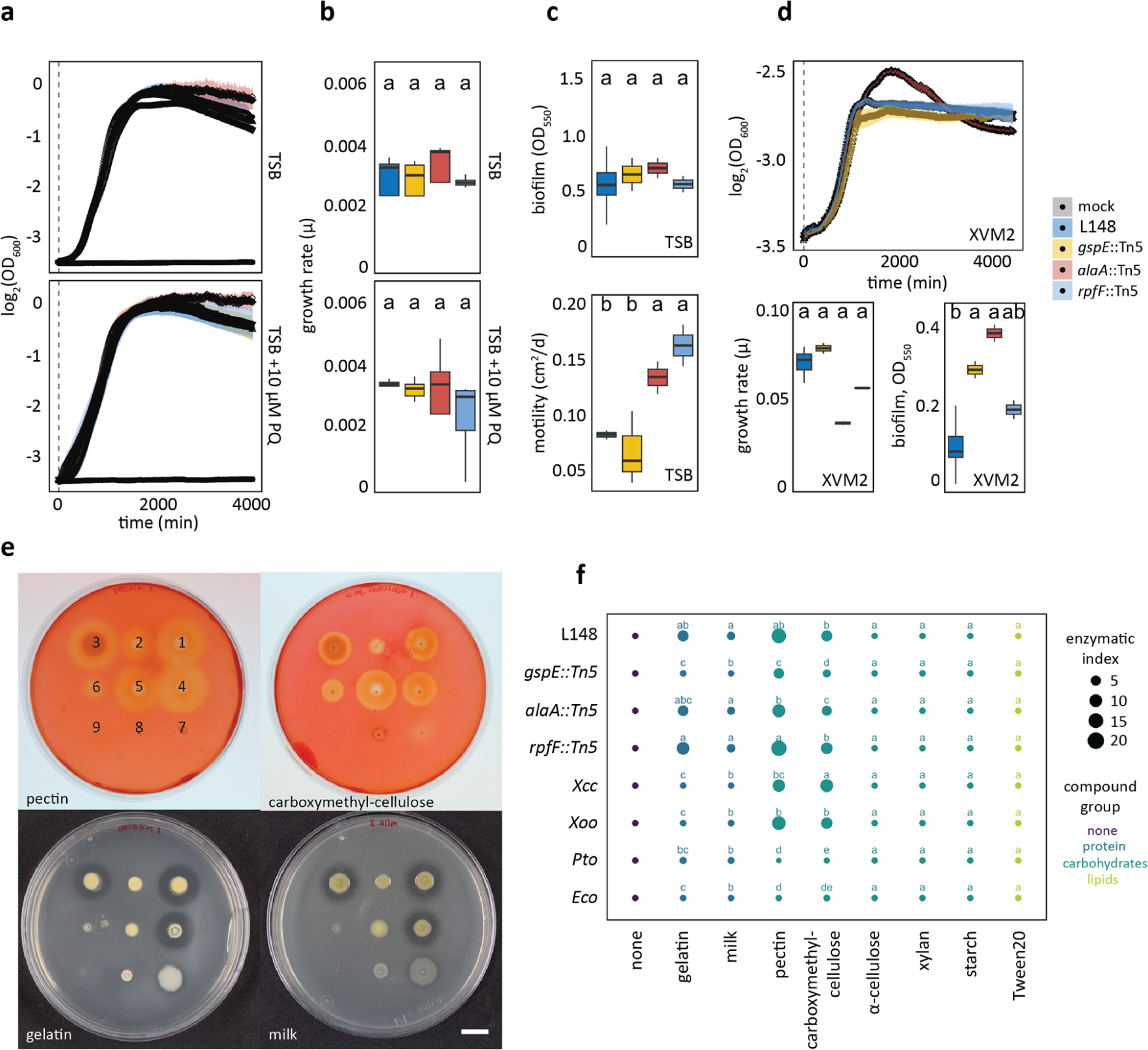
The *Xanthomonas* L148 *gspE*::Tn5 mutant exhibits compromised extracellular secretion activity. **a,b**, Growth curves (**a**) and rates (**b**) of *Xanthomonas* L148::Tn5 candidate mutant strains in TSB upon chronic exposure to 0 or 10 µM PQ for 4000 min (2 independent experiments each with 3 biological replicates). **c**, Biofilm production and motility of *Xanthomonas* L148::Tn5 candidate mutant strains in TSB medium (2 independent experiments each with 2–3 biological replicates). **d**, Growth curves, growth rates, and biofilm production of *Xanthomonas* L148::Tn5 candidate mutants in XVM2 (2 independent experiments each with 2–3 biological replicates). **e**, Exemplary images of plate assays for secretion activities of bacterial strains (1 = wildtype *Xanthomonas* L148; 2 = *gspE*::Tn5; 3 = *alaA*::Tn5; 4 = *rpfF*::Tn5; 5 = *Xanthomonas campestris* pv*. campestris* [*Xcc*]; 6 = *X. oryzae* pv*. oryzae* [*Xoo*]; 7 = *P. syringae* pv. *tomato* DC3000 [*Pto*]; 8 = *E. coli* HB101 [*Eco*]; and 9 = mock) for the carbohydrates pectin and carboxymethylcellulose, and gelatin and milk proteins. **f**, Enzymatic indices for bacterial strains grown on TSB supplemented with 0.1% substrates (proteins: gelatin and milk; carbohydrates: pectin, carboxymethyl-cellulose, α-cellulose, xylan, and starch; lipids: Tween20) after 2 day-incubation at 28 °C (3 biological replicates). The enzymatic indices were calculated by subtracting the size of the colony with the zone of clearance, indicative of substrate degradation by the strain after 2–3 d. **b,d**, the growth rate, μ, was calculated by running rolling regression with a window of 5 h along the growth curves to determine the maximum slope. **b–d, f**, Different letters indicate statistically significant differences (ANOVA with *post hoc* Tukey’s test, *P* ≤ 0.05). Results in **b**, **c** and **d** are depicted as box plots with the boxes spanning the interquartile range (IQR, 25^th^ to 75^th^ percentiles), the mid-line indicates the median, and the whiskers cover the minimum and maximum values not extending beyond 1.5x of the IQR.

To gain insight into the evolution of the pathogenicity of *Xanthomonas* L148, available genomes of other Xanthomonadales members, including the potentially pathogenic close-relative *Xanthomonas* L131^17^ and *Xanthomonas* L70 in the AtSPHERE^25^, together with several *Xanthomonas* pathogens and *X. massiliensis*, an isolate from human feces were interrogated for the occurrence of secretion systems and their potential CAZyme catalogues. In general, all Xanthomonadales strains encode both T1SS and T2SS genes (Supplemental Figure S8a). The pathogenic and potentially pathogenic Xanthomonadales strains have expanded their CAZyme repertoire with proclivities for plant cell wall components: α-, β-glucans, β-mannans, arabinan, cellulose, and pectin (Supplemental Figure S8b-c). This indicates that though secretion systems are prevalent among the Xanthomonadales members, CAZyme repertoire expansion might be key feature of pathogenic and potentially pathogenic strains.

Because of the *in planta, ex planta*, and *in vitro* phenotypes, we focused on *gspE*::Tn5 mutant and characterized it extensively. To establish that *gspE* determines *rbohD*-dependent pathogenicity, we generated two independent *gspE* deletion mutant strains (*ΔgspE*_1 and *ΔgspE*_2) via homologous recombination. Both of the *gspE* deletion mutants as well as the *gspE*::Tn5 mutant showed loss of secretion activities and failed to cause disease in *rbohD* plants (Figure 6a-b). Taken together, *gspE*, an integral component of T2SS, is essential for *Xanthomonas* L148 pathogenicity on *rbohD* mutant plants.

**Figure 6.**
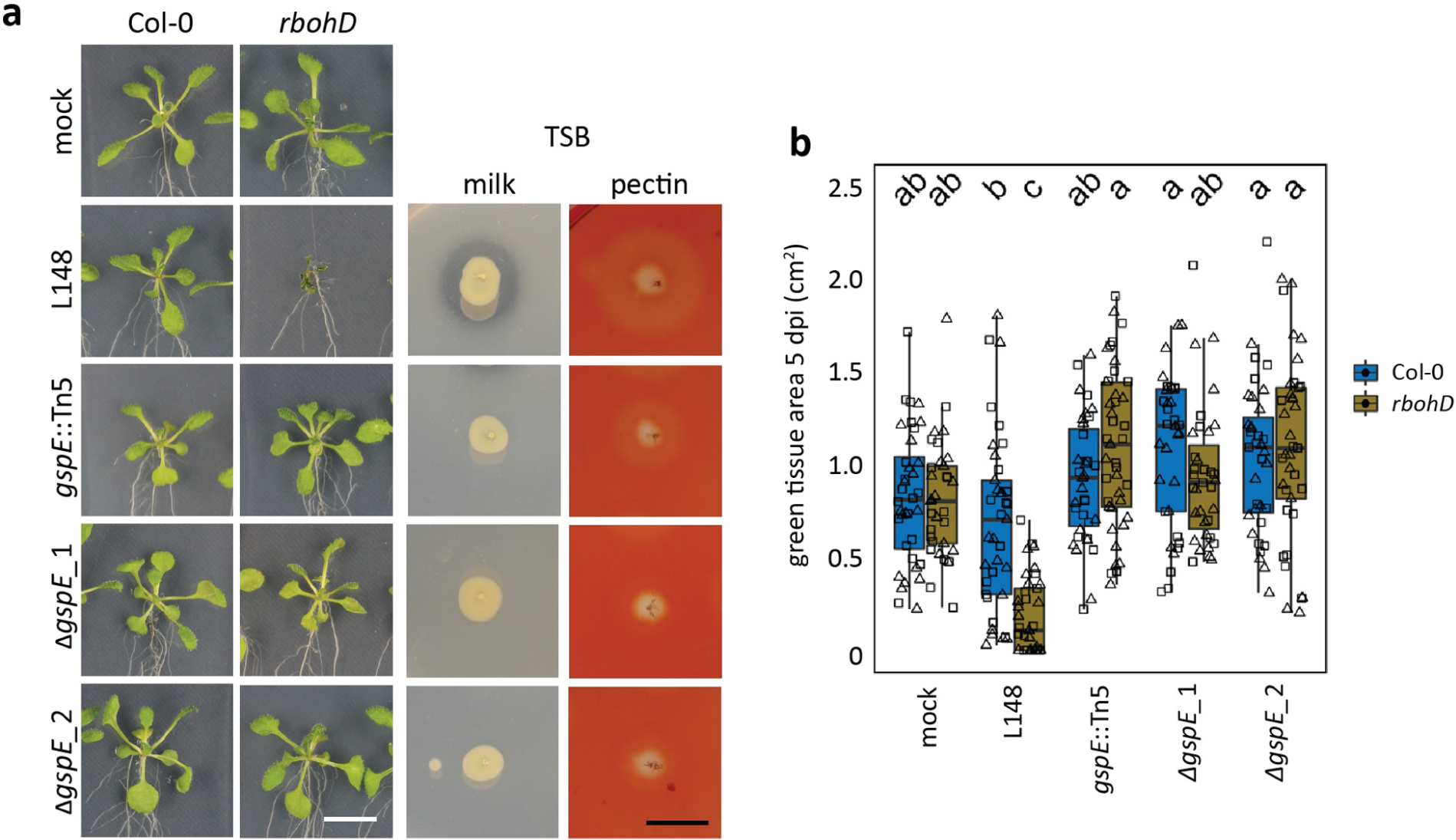
The T2SS component *gspE* is a genetic determinant for the loss of the *rbohd*-dependent pathogenicity of *Xanthomonas* L148. **a**, Images of axenic Col-0 and *rbohD* plants flood-inoculated (OD600=0.005) with the wildtype *Xanthomonas* L148, *gspE*::Tn5 mutant and 2 *ΔgspE* lines at 5 dpi. Plate images of secretion activities of the bacterial strains grown on TSB with either milk or pectin as substrate after 2–3 days. **b**, Measured green tissue area as the plant health parameter for Col-0 and *rbohD* plants flood-inoculated with the bacterial strains at 5 dpi (2 independent experiments each with 3–5 biological replicates). Different letters indicate statistically significant differences (ANOVA with *post hoc* Tukey’s test, *P* ≤ 0.05). Results in **b** are depicted as box plots with the boxes spanning the interquartile range (IQR, 25^th^ to 75^th^ percentiles), the mid-line indicates the median, and the whiskers cover the minimum and maximum values not extending beyond 1.5x of the IQR.

### Plant ROS suppresses T2SS genes including *gspE* of *Xanthomonas* L148

*Xanthomonas* L148 pathogenicity is exerted in the absence of ROS through *RBOHD*, while our *in vitro* results do not indicate general cellular toxicity of ROS. Thus, it can be assumed that *RBOHD*-mediated ROS production suppresses virulence of *Xanthomonas* L148. To gain insight into this, we conducted *in planta Xanthomonas* L148 bacterial transcriptome profiling for Col-0 and *rbohD* plants^26^. Plants were flood-inoculated with *Xanthomonas* L148 and shoots were sampled at 2 dpi, a time point at which bacterial titers were still indistinguishable; these later became significantly different between Col-0 and *rbohD* leaves at 3 dpi (Figure 2f). Thus, with the bacterial transcriptomes observed at this time point, one can exclude the possibility that the differences in expression are due to the different bacterial population densities known to affect bacterial transcriptome^31^.

Principal component (PC) analysis revealed that *in planta Xanthomonas* L148 transcriptomes were distinct in Col-0 and *rbohD* plants (Figure 7b). Statistical analysis revealed 2,946 differentially expressed genes (DEGs) upon comparing *in planta* bacterial transcriptomes in Col-0 with *rbohD* leaves (threshold: q-values < 0.05): 563 genes were up-regulated and 2,474 genes were down-regulated in Col-0 compared with *rbohD* plants (Figure 7a and c, See Supplementary Dataset S2 for the details on DEGs). Strikingly, most T2SS apparatus genes including *gspE* were down-regulated in Col-0 as compared to *rbohD* (Figure 7c–e). The DEGs were significantly enriched for the candidate genes detected from the *Xanthomonas* L148::Tn5 mutant screening (29 up-regulated and 73 down-regulated out of 214 genes in Col-0 as compared to *rbohD*-inoculated plants, hypergeometric test, p-value = 1.00E-10***), which highlights a remarkable concurrence of the genetic evidence with the bacterial transcriptome profiles obtained *in planta* (Figure 7a). The DEGs were also significantly over-represented for carbohydrate-active enzymes (CAZyme, 4 up-regulated, 49 down-regulated out of 135 in Col-0 as compared to *rbohD*-colonized plants, hypergeometric test, p-value = 1.53E-12***, Figure 7a, c), which is consistent with the notion that CAZymes function in virulence. Moreover, six *Xanthomonas* L148::Tn5 mutants have an insertion in genes annotated as CAZymes, five of which are significantly down-regulated in Col-0 as compared with *rbohD* inoculated plants. The significantly down-regulated CAZymes in Col-0 plants can potentially degrade plant cell wall components cellulose, pectin, α-glucan, β-glucan, and β-mannan (Figure 7c, Supplementary Figure S9). Pathway enrichment analysis revealed that upregulated gene clusters such as clusters 3, 9, and 14 are enriched for biological functions related to chemotaxis and attachment (K15125, K13924, and K05874), while gene clusters down-regulated in Col-0 such as clusters 8, 10, and 12 are enriched for pathways involved in transport and detoxification processes (K02014 and K00799, Figure 7f, See Supplementary Dataset S3 for the clustering membership and the enriched GO terms).

**Figure 7.**
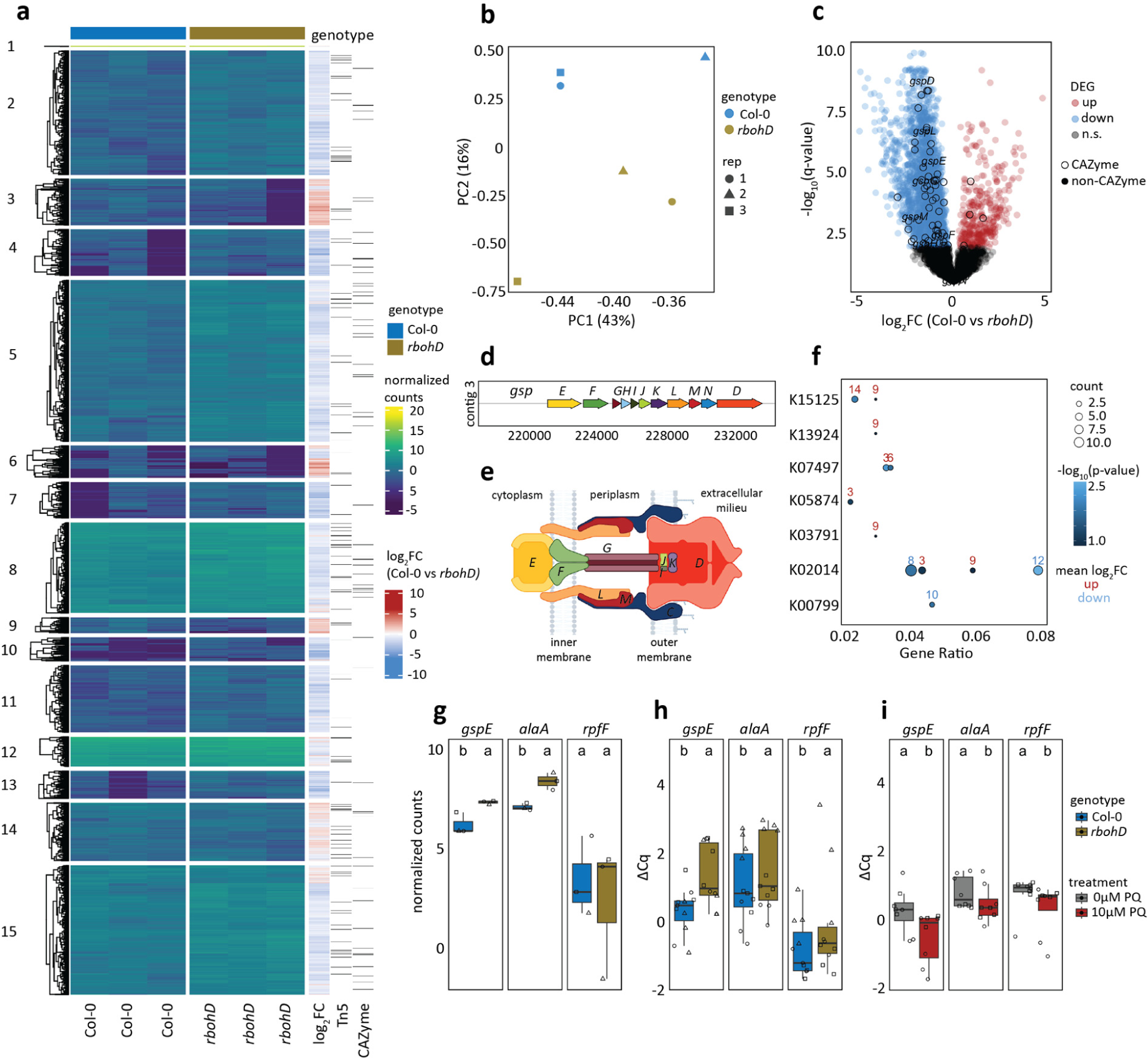
Plant ROS suppress T2SS genes including *gspE* of *Xanthomonas* L148. **a**, Heatmap representation of *in planta* bacterial transcriptome landscape of the wildtype *Xanthomonas* L148 in Col-0 and *rbohD* plants. Leaves of 2-week-old plants were flood-inoculated with L148 and samples were taken at 2 dpi. Gene clusters were based on k-means clustering of the normalized read counts. DEGs were defined based on q-value < 0.05. Sidebars indicate the log2 fold changes of Col-0 compared with *rbohD*, *Xanthomonas* L148::Tn5 candidate genes (the 214 candidates), and the genes annotated as CAZyme. **b**, Principal component (PC) analysis of the *in planta Xanthomonas* L148 transcriptome for DEGs in Col-0 and *rbohD* plants. **c**, Volcano plot of the DEGs with which T2SS component genes were labelled and CAZymes highlighted. **d**, Genomic architecture of the T2SS genes. **e**, Graphical representation of T2SS assembly. **f**, KEGG pathway enrichment analysis of the gene clusters (indicated in numbers) in **a**. **g**, RNA-Seq normalized counts of *gspE*, *alaA*, and *rpfF*. **h**, Independent qRT-PCR experiments for *in planta* expression profiling of *gspE*, *alaA*, and *rpfF*. Experiments were performed as in RNA-seq with 2 independent experiments each with 3–4 biological replicates. **i**, qRT-PCR *in vitro* expression profiling of *gspE*, *alaA*, and *rpfF* in *Xanthomonas* L148 wildtype strain grown in TSB ± 10 μM PQ for 24 h (2 independent experiments each with 3–4 biological replicates). **h,i**, Gene expression was normalized against the housekeeping gene *gyrA*. Different letters indicate statistically significant differences (ANOVA with *post hoc* Tukey’s test, *P* ≤ 0.05). Results in **g–i** are depicted as box plots with the boxes spanning the interquartile range (IQR, 25^th^ to 75^th^ percentiles), the mid-line indicates the median, and the whiskers cover the minimum and maximum values not extending beyond 1.5x of the IQR. Some illustrations were created with BioRender.

Upon closer inspection, expression of the identified candidate genes *gspE* and *alaA* was strongly repressed while *rpfF* was marginally downregulated in Col-0 compared to *rbohD*, which supports the hypothesis that these genes are required and thus tightly regulated by immunocompetent wild-type Col-0 plants to prevent disease progression (Figure 7g). These findings were re-confirmed in independent experiments using qRT-PCR where all the candidate genes were suppressed in Col-0 compared to *rbohD* plants (Figure 7h). It can be postulated that ROS directly regulates the expression of these genes. Therefore, *Xanthomonas* L148 bacterial cells were grown *in vitro* in the presence of PQ followed by gene expression analysis. We found that the expression of the candidate genes *gspE, alaA*, and *rpfF* is suppressed in *Xanthomonas* L148 upon chronic exposure to ROS (Figure 7i). Taken together, these findings suggest that *Xanthomonas* L148 colonization triggers RBOHD-mediated ROS production, which directly inhibits the expression of genes related to virulence, in particular components of the T2SS on Col-0 plants. By contrast, the absence of ROS production in *rbohD* mutant plants switches on the pathogenicity of *Xanthomonas* L148, leading to disease onset.

### RBOHD-mediated ROS turns *Xanthomonas* L148 into a beneficial bacterium

The phyllosphere microbiota are known to confer protection against foliar pathogens^32^ and thus even a conditionally pathogenic microbiota member may provide beneficial services to its plant host. To address this question, Col-0 and *rbohD* plants were pre-colonized with wild-type *Xanthomonas* L148 or *gspE*::Tn5 mutant strain for five days and were then challenged with the bacterial pathogen *Pto*. Bacterial titers of *Xanthomonas* L148 and *Pto* were determined for the endophytic and total leaf compartments at 0 and 3 dpi. As *Xanthomonas* L148 killed *rbohD* mutant plants, we were not able to measure *Pto* titers under this condition. Pre-colonized Col-0 plants with either the wild-type *Xanthomonas* L148 or *gspE*::Tn5 mutant had increased resistance against *Pto* (Figure 8a-c). Interestingly, *rbohD* mutant plants pre-colonized with *gspE*::Tn5 strain showed increased resistance against *Pto*, resembling *Xanthomonas* L148 pre-colonized Col-0 plants (Figure 8a, c). Further, Col-0 and *rbohD* plants pre-colonized with *gspE*::Tn5 had slightly better plant performance than the non-inoculated plants after *Pto* challenge (Supplementary Figure S10a-b), suggesting that the strain promotes plant fitness in the presence of pathogens. Invasion by *Pto* did not result in a significant decline in *Xanthomonas* L148 and *gspE*::Tn5 populations (Figure 8b), indicating a strong colonization competence and resistance of the commensal *Xanthomonas* L148 against pathogen invasion. In summary, these results revealed that RBOHD-produced ROS turns the potentially harmful *Xanthomonas* L148 into a beneficial bacterium, thereby protecting the plant from aggressive pathogen colonization.

**Figure 8.**
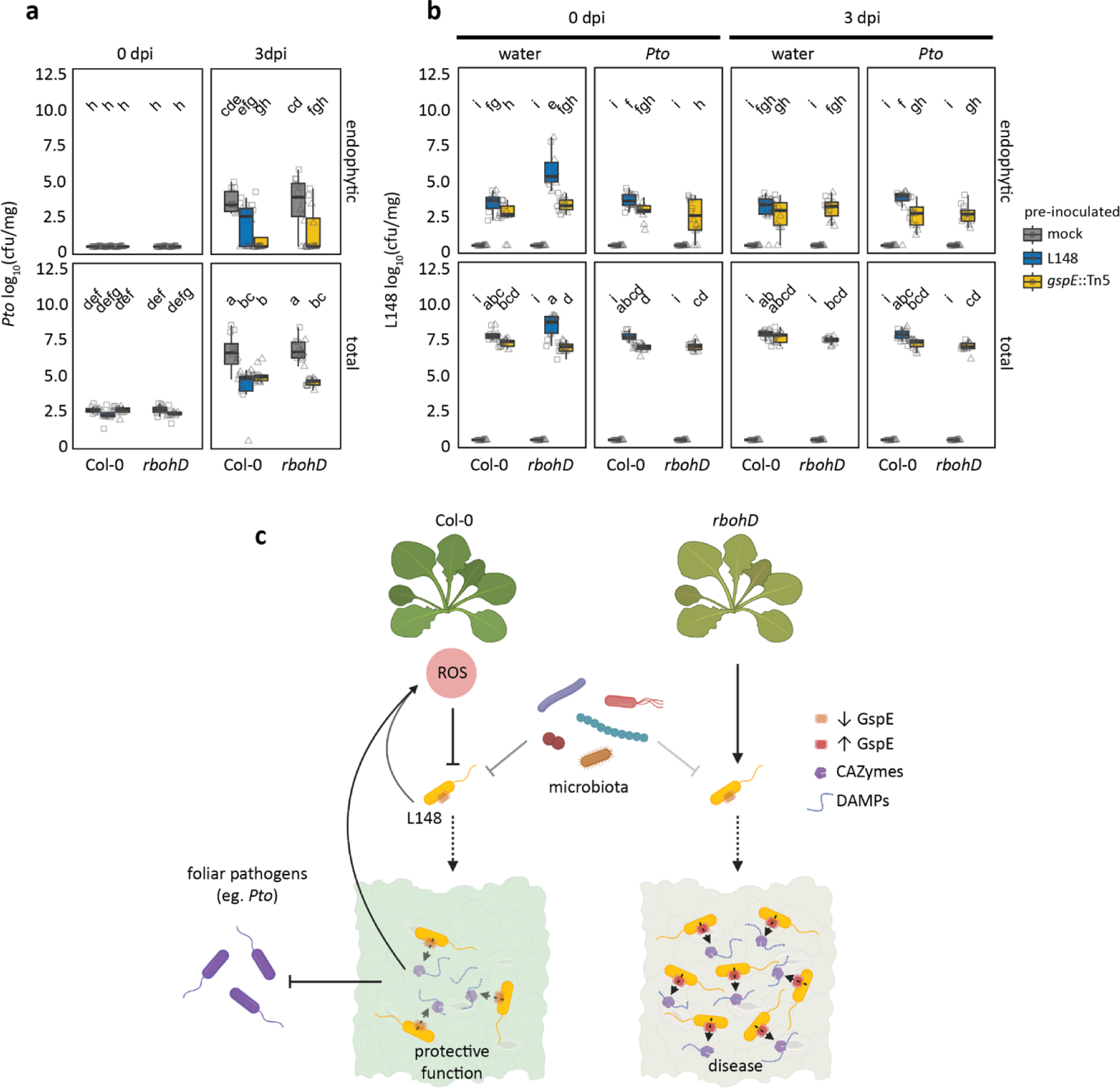
RBOHD-mediated ROS turn *Xanthomonas* L148 into a beneficial bacterium. **a, b**, 14-day-old Col-0 and *rbohD* plants grown on agar plates were flood-inoculated with wildtype *Xanthomonas* L148 and *gspE*::Tn5 (OD600=0.005) for 5 days followed by spray infection with *Pto*. Bacterial titers were determined at 0 and 3 dpi (**a**, *Pto*; **b**, L148) (2 independent experiments each with 6 (**a**) or 3–5 (**b**) biological replicates). Different letters indicate statistically significant differences (ANOVA with *post hoc* Tukey’s test, *P* ≤ 0.05). Results in **a**–**b** are depicted as box plots with the boxes spanning the interquartile range (IQR, 25^th^ to 75^th^ percentiles), the mid-line indicates the median, and the whiskers cover the minimum and maximum values not extending beyond 1.5x of the IQR. **c**, Mechanistic model for plant ROS licensing of co-habitation with a potentially pathogenic *Xanthomonas* L148 commensal, where the microbe releases MAMPs that are perceived by plants and trigger ROS production. The T2SS delivers CAZymes to the host to degrade cell wall liberating DAMPs and/or the CAZymes act as a MAMP, which both can potentially bolster ROS generation. The ROS then acts as a molecular beacon for *Xanthomonas* L148 to suppress its pathogenicity, in particular by dampening the activity of T2SS resulting in a negative feedback regulation of the bacterial activity by the plant host. We propose that in wild-type Col-0 plants, the ROS-and the microbiota-mediated suppression of *Xanthomonas* L148 promotes the cooperative behavior of L148 with the host plant and in turn confers protective function against subsequent invasion by foliar pathogens. In the case of *rbohD* mutant plants wherein plant ROS is absent, *Xanthomonas* L148 virulence is unlocked, resulting in disease. Some illustrations were created with BioRender.

## Discussion

Despite extensive studies on how plants recognize microbes and transduce signals within the plant, how immune outputs control the growth and behavior of microbes is still largely unknown. Furthermore, we mostly lack a mechanistic explanation for why certain microbes are sensitive to particular immune responses. In this study, we have investigated the impact of the RBOHD-mediated ROS burst as an early immune output on 20 taxonomically diverse bacteria isolated from healthy *A. thaliana* plants and demonstrated the poor association between RBOHD-mediated ROS burst and bacterial colonization (Supplementary Figure S4). This highlights the notion that the perception of the microbial signal, followed by the cascade of immune signals, and immune execution leading to the restriction of microbial colonization are distinct events. This corroborates our previous finding that plant and bacterial transcriptome responses are largely uncoupled during an early stage of infection^26^. In this study, we have revealed a mechanism in which RBOHD-mediated ROS changes the growth and behavior of a leaf commensal, *Xanthomonas* L148. This is a significant advance in our understanding of how plant immune responses manipulate bacterial growth and behavior.

We have demonstrated that plant ROS licenses co-habitation with a potentially detrimental *Xanthomonas* L148 while it trains L148 to guard against aggressive leaf pathogens. Our results show that the plant host constrains proliferation of this microbiota member by means of ROS as a molecular message to harness it for its own benefits. Ecological and reductionist studies have revealed that potentially pathogenic strains populate plant hosts without causing disease, and these strains are considered as *bona fide* constituents of the plant microbiota^29, 33–35^. Some of these potentially detrimental strains can be deleterious to the host in mono-associations^29, 33–37^. However, the adverse effects of these potentially pathogenic microbes depend on the host, the environment, and the co-occurring microbes^17, 32, 34–38^. It has been shown that simultaneous defects in PTI and the vesicle trafficking pathway under high humidity led to dysbiosis in the phyllosphere and plant disease^11–12^. It appears to be a universal pattern across multicellular organisms that ROS modulates the structure, composition, and function of microbiota. In mice, a decrease in mitochondria-derived ROS is associated with increased gut microbiota diversity^39^. Also, ROS produced via the NOX1 pathway in the colon drives anaerobic growth of *Citrobacter rodentium* and in turn remodel the epithelial milieu^40^. In plants, ROS induces the phytohormone auxin secretion by a beneficial rhizobacterium *Bacillus velezensis* to protect against the damaging effects of plant-derived ROS, allowing efficient root colonization of *B. velezensis*^18^. ROS production in roots constrains *Pseudomonas* establishment in the rhizosphere^16^. It has also been genetically shown that RBOHD-mediated ROS production is integral for maintaining leaf microbiota homeostasis by keeping potentially harmful bacterial members at bay^17^. Nevertheless, the mechanisms by which the plant host selectively constrains potentially pathogenic members of the microbiota and whether these strains are functional to their host remains unclear. Here, through a bacterial genome-wide transposon mutant screen and *in planta* transcriptomics, we have revealed that plant ROS acts as a signaling cue for the potentially pathogenic commensal *Xanthomonas* L148 to suppress its virulence by downregulating its T2SS while promoting its beneficial function.

Other members of the phyllosphere microbiota may partially contribute to attenuating the deleterious effects of *Xanthomonas* L148. However, *RBOHD* is needed for full suppression of L148 deleterious activity in the community context (Supplementary Figure S6), which is consistent with the observation that *Xanthomonas* L131, a closely-related strain of L148, exerts its detrimental impact on *rbohD* mutant plants in a community context^17^. Closely related, innocuous strains of the plant microbiota out-compete or antagonize its potentially pathogenic counterparts, thereby preventing disease progression but enabling the persistence and co-existence of these strains in nature^23, 39^. However, this phenomenon is accession and strain-specific as this commensal-mediated protection is lost in some plant genotypes and a particular harmful *Pseudomonas* strain predominates the microbial community^39^. Thus, allowing potentially pathogenic strains within the microbiota requires stringent control of their function and behavior by host immunity sectors and is facilitated in parts by other members of the plant microbiota.

We have demonstrated that the pathogenicity of *Xanthomonas* L148 depends on the T2SS component *gspE* (Figure 3d-f, 6a-b). The loss of the killing effect of *gspE*::Tn5 mutants strains on *rbohD* mutant plants can be explained by its compromised secretion activities and hampered colonization of *rbohD* leaves (Figure 4b, 5e-f, 6a-b, Figure 7c, and Supplementary Figure 9a-b). The T2SS is often utilized by plant pathogens to deliver CAZymes which degrade plant cell walls, allowing host invasion and promoting disease^30^. For instance, the T2SS allows the root commensal *Dyella japonica* MF79 to efficiently colonize the host and is required for virulence of pathogenic *Dickeya dadantii*^24, 41^. Secreted CAZymes could also trigger immune responses such as ROS burst via direct recognition of the CAZyme as a MAMP or release of recognized plant-derived Damage Associated Molecular Patterns (DAMPs) due to their enzymatic action^41–44^. Indeed, we have shown that live T2SS-deficient *gspE*::Tn5 L148 mutant elicited less ROS than wild-type L148, whereas heat-killed wild-type L148 and *gspE*::Tn5 mutant elicited undistinguishable ROS burst, implying that T2SS-mediated CAZyme secretion may further enhance the ROS response (Figure 4c). Plant ROS might act as a counter-defense of L148 invasion via CAZymes by dampening T2SS expression (Figure 7c, Supplementary Figure 9a-b). Considering that wild-type *Xanthomonas* L148 and the *gspE*::Tn5 mutant had similar leaf colonization patterns in wild-type Col-0 plants (Figure 4b), this counter-defense likely functions to attenuate T2SS activity and make *Xanthomonas* L148 a commensal bacterium in wild-type Col-0 plants. Thus, we propose a model according to which the interaction of *Xanthomonas* L148 and Col-0 plants is based on a delicate balance driven by host ROS levels, resulting in a negative feedback loop to control the potentially pathogenic commensal (Figure 8c). Moreover, the plant protective function of *Xanthomonas* L148 against the pathogen *Pto* is not genetically coupled with its *gspE*-dependent pathogenic potential, as the *gspE*::Tn5 mutant can still confer significant resistance against *Pto* in both Col-0 and *rbohD* mutant plants (Figure 8a, Supplementary Figure S10a-b). These findings suggest an important role of the T2SS in the establishment of microorganisms in host tissues, making it conceivable that it is targeted by the host to manipulate microbial behavior. Our finding that RBOHD-mediated ROS targets *Xanthomonas* T2SS provides a new mechanism and concept that plant immunity surveils potentially detrimental members of the plant microbiota by suppressing the T2SS via ROS.

We have revealed different ROS burst patterns in response to individual members of the plant microbiota that can be categorized into three classes of immune reactivity: immune-active strains can elicit ROS with intact cells; immune-evasive strains only induce ROS when they are heat-killed; and immune-quiescent strains do not elicit ROS whether alive or dead (Figure 1c,d and Supplementary Figure S2). Immune-evasive strains can possibly conceal their detection by preventing MAMP release or secrete proteins that degrade/sequester self-derived MAMPs or that target host immune components to suppress immune activation^46^.

We have observed that most microbiota members of *A. thaliana* are perceived through the surface-resident PRR EFR (Supplementary Figure S2), indicating that EF-Tu peptides serve as major bacterial molecules eliciting defense programs in our experimental setup. Consistent with this, a number of strains increased colonization in *efr* mutant plants compared to wild-type Col-0, which emphasizes the fundamental link of microbial perception with bacterial colonization (Supplementary Figure S3). Our observation also coincides with a GWAS study in which *EFR* was found as a plausible genetic determinant of responses to varying MAMP epitopes in natural populations of *A. thaliana*^47^. Although EFR is a Brassicaceae lineage-specific innovation^7^, other EF-Tu fragments seem to be recognized by yet-unknown receptors and are immunogenic to some rice cultivars^48^. Also, interfamily transfer of *A. thaliana* EFR to solanaceous species is sufficient to confer broad-spectrum resistance to pathogens, indicating that components acting downstream of EFR perception are at least in parts evolutionarily conserved^4^. These findings suggest that EF-Tu peptides might be a prevalent microbial motif for host detection in various plant species.

Emerging evidence suggests that plant immunity modulates microbial processes required for virulence in addition to its effects on general microbial metabolism, including protein translation^31, 45^. For instance, the secreted aspartic protease SAP1 inhibits *Pto* growth by cleaving the *Pto* protein MucD in *A. thaliana* leaves^50^. Plants target the iron acquisition system of *Pto* to inhibit *Pto* growth during effector-triggered immunity^31^. The defense phytohormone salicylic acid and the specialized metabolite sulforaphane inhibit the type III secretion system of pathogenic *Pto*^45, 51^. Our finding is consistent with the notion that plant immunity targets microbial virulence to allow microbes to cohabit, which can be a better plant strategy than eliminating microbes as plants need to maintain a functional microbiota and potentially harmful microbes can even provide a service to the host.

## Materials and Methods

### Plant materials and growth conditions

The *A. thaliana* Col-0 accession was the wild-type and the genetic background of all the mutants utilized in this study. The mutants *fls2*^7^ (*SAIL_691C4*), *efr*^8^ (SALK_068675), *cerk1*^10^ (GABI_096F09), *fec*^11^, *bbc*^11^, and *rbohD*^13^ (*atrbohD D3*) were previously described. For agar plate assays, seeds were sterilized with Cl_2_ gas for 2 h^52^. Seeds were then stratified for 2–3 days at 4 °C on 0.5x Murashige and Skoogs (MS) medium agar with 1% sucrose, germinated for 5 days, and subsequently transplanted to 0.5x MS plates without sucrose. Plants were grown in a chamber at 23 °C/23 °C (day/night) with 10 h of light. Then, 14-day-old seedlings were inoculated with bacterial strains and were harvested or phenotyped at the indicated time points. For ROS burst and infiltration patho-assays, plants were grown in greenhouse soil for 5–6 weeks in a chamber at 23 °C/23 °C (day/night) with 10 h of light and 60% relative humidity (See Supplementary Table S1 for details of the plant genotypes used).

### Bacterial strains and growth conditions

All the bacterial strains derived from the AtSPHERE were previously described^25^. *Pseudomonas syringae* pv. *tomato* DC3000 (*Pto*) and *Pto* lux were described previously^53–54^. All bacterial strains were grown in 0.5x Tryptic Soy Broth (TSB) for 24 h, harvested through centrifugation, washed twice with sterile water, and diluted to the appropriate OD_600_ (See Supplementary Table S2 for the list of bacterial strains used).

### ROS burst measurement

ROS burst was determined as in Smith and Heese, 2014 with slight modifications^55^. In brief, bacterial strains were grown in TSB at 28 °C for 16–18 h with shaking at 200 rpm. Cells were harvested, washed twice with sterile water, and diluted to OD_600_=0.5 in sterile water. The day before the assay, leaf discs (4 mm) from leaves of the same physiological state and size from 5-to-6-week-old plants grown in a chamber at 23 °C/23 °C (day/night) with 10 h of light were harvested, washed twice with sterile water every 30 min, immersed in sterile water in 96-well plates, and incubated in the same growth chamber for 20 h. Prior to the assay, the elicitation solution was prepared by adding 5 µL 500x horseradish peroxidase (HRP, P6782-10MG, Sigma-Aldrich) and 5 µL 500x luminol (A8511-5G, Sigma-Aldrich) to 2.5 mL of bacterial suspension, 1 µM MAMP solutions (flg22 [ZBiolab inc.], elf18 [Eurofins], chitinDP7 [N-acetylchitoheptaose, GN7, Elicityl]), or sterile nanopure water as mock. During the assay, the water was carefully removed from the 96 well-plate and 100 µL of the elicitation solution was added to the 96-well plate. With minimal delay, the luminescence readings were obtained for 60 min using a luminometer (TrisStar2 Multimode Reader, Berthold).

### Commensal bacterial colonization assay

To prepare the bacterial inoculum, all bacterial strains were grown in 0.5x TSB for 24 h, harvested through centrifugation, washed twice with sterile water, and suspended in sterile water (final OD_600_=0.005). Two-week-old seedlings grown on 0.5x MS medium agar in a chamber at 23 °C/23 °C (day/night) with 10 h of light were flood-inoculated with these bacterial suspensions and incubated in the same growth chamber. Leaf samples were aseptically harvested at 3 to 5 dpi, weighed, and plated for two compartments: for the total compartment, leaves were directly homogenized in 10 mM MgCl_2_ with a homogenizer (TissueLyser III, Qiagen), serially diluted with 10 mM MgCl_2_, and plated on 0.5x TSB agar; for the endophytic compartment, leaves were surface-sterilized with 70% ethanol for 1 min, washed twice with sterile water, homogenized, serially diluted, and then plated as for the total compartment. Colonies were allowed to grow at 28 °C, and photographs were taken for 1 to 3 days. Colonization was expressed as cfu mg^-1^ sample.

### Generation of bacterial mutants

A *Xanthomonas* L148::Tn5 library was constructed via conjugation of *Xanthomonas* L148 with *E. coli* SM10*λpir* harboring puTn5TmKm2^56^ in which both strains were mixed in equal portions (OD_600_=0.10), spot-plated on TSB medium, and incubated for 2 d at 28 °C. The resulting mating plaques were diluted and plated on TSB with kanamycin and nitrofurantoin for selection for L148 transformants and counter-selection against *E. coli*, respectively. To constitute the entire library, around 7,000 individual colonies were picked, re-grown in 0.5x TSB, aliquoted for glycerol stocks, and stored at −80 °C. Around 20 strains from this *Xanthomonas* L148::Tn5 library were randomly selected for confirmation of Tn5 insertion in the genome via nested PCR (first PCR with primers FDE117 and FDE118; second PCR with primers FDE119 and mTn5AC) and the final amplicons were Sanger-sequenced (see Supplementary Table S3 for details). For the generation of targeted deletion mutants for *gspE*, the pK18mobsacB suicide plasmid^57^ (GenBank accession: FJ437239) was PCR linearized (primers FDE234 and FDE235) with Phusion Taq polymerase (F-5305, Thermo Scientific); 750 bp of upstream (primers FDE278 and FDE279) and downstream (primers FDE280 and FDE281) flanking regions of *gspE* coding sequence with terminal sequences overlapping with the linearized pK18mobsacB were amplified using Phusion Taq polymerase (F-5305, ThermoScientific) and were sequence-verified. The plasmid construct was assembled using Gibson cloning^58^ (E5510S, New England Biolabs) following the manufacturer’s instructions. The plasmid construct was transformed into *E. coli* cells (DH5⍺ strain) and then delivered into *Xanthomonas* L148 via triparental mating with the helper strain pRK600^59^. Transformants were selected using kanamycin and nitrofurantoin and the second homologous recombination was induced with sucrose in 0.5x TSB. The deletion mutants were individually picked and stored at −80 °C in glycerol stocks and were verified by PCRs (using primers FDE196 and FDE197 for the presence of the plasmid with the inserts; primers FDE125 and FDE126 for the presence of *gspE* gene in the genome; and primers FDE279 and FDE280 for the removal of *gspE* gene in the genome) and Sanger-sequencing, and were plated on 0.5x TSB containing 10 μg/mL kanamycin. True deletion mutants should not contain the plasmid, lose the *gspE* gene, and be sensitive to kanamycin (See Supplementary Table S3 for list of primers and PCR profile used).

### L148::Tn5 library 96-well screening

Seedlings of *rbohD* were grown in 96-well plates with 0.5x MS agar with 1% sucrose for 14 days. Concomitantly, the Tn5 insertion mutants (∼7,000 individually picked colonies) were grown in 96-well plates with TSB at 28 °C for 3 days with 200 rpm agitation till saturation. The resulting bacterial suspension was diluted six times (resulting in a concentration of approximately 6×10^9^ bacterial cells per mL) and 20 µL aliquots were inoculated onto the seedlings. Plants were phenotyped for survival after 5 days. The resulting 214 *Xanthomonas* L148::Tn5 candidate strains which showed the loss of the *rbohD* killing activity from the two independent 96-well plate screenings were genotyped to identify the Tn5 insertion locus in the genome via nested PCR (first PCR with primers FDE117 and FDE118; second PCR with primers FDE119 and mTn5AC) and the final amplicons were Sanger-sequenced (see Supplementary Table S3 for list of primers and PCR profile used and Supplementary Figure S7). The 124 *Xanthomonas* L148::Tn5 candidate mutants which have insertions on genes with functional annotations (please see Supplementary Dataset S1 for the list) were further screened using plants grown in agar plates to re-evaluate the phenotypes as described for the commensal bacterial colonization assay.

### In vitro assays

For instantaneous ROS treatment, *Xanthomonas* L148 was grown for 24 h, pelleted, and diluted to OD_600_ = 0.02. A 500 µL of the bacterial suspension was mixed with H_2_O_2_ (H10009-500ML, Sigma-Aldrich) at final concentrations of 0–2000 µM, incubated for 30 min, and plated for colony counts. Similarly, 500 μL of the bacterial suspension was mixed with 1 mM xanthine (X7375-10G, Sigma-Aldrich) and 10 U/mL xanthine oxidase from bovine milk (X4875-10UN, Sigma-Aldrich) to generate O_2-1_, and samples were plated at different time points (1 mol of xanthine is converted to 1 mol O_2-1_ with 1 U xanthine oxidase at pH 7.5 at 25 °C in a min, thus 0, 2, 4, 10, 20, 40, 60, and 80 min incubations should have produced O_2-1_ equivalent to 0, 50, 100, 250, 500, 1000, 2000 µM respectively) for colony counts. Chronic exposure to ROS was implemented by growing the strains in TSB ± 10 µM paraquat (856177-1G, Sigma-Aldrich), a ROS-generating compound, for three days while obtaining OD_600_ readings using spectrophotometer (Tecan Infinite Microplate reader M200 Pro) to calculate growth curves and rates. The candidate *Xanthomonas* L148::Tn5 mutants were phenotyped *in vitro* via growing bacterial culture with an initial inoculum of 10 µL OD_600_=0.1 in 96-well plates supplemented with 140 µL TSB or XVM2 (a minimal medium designed for *Xanthomonas* strains^60^) for three days while obtaining absorbance readings at OD_600_ using a spectrophotometer (Tecan Infinite Microplate reader M200 Pro) to calculate growth curves and rates. The resulting cultures were gently and briefly washed with water and cells adhering on the plates were stained with 0.1% crystal violet (27335.01, Serva) for 15 min. The staining was solubilized with 125 μL 30% acetic acid (A6283, Sigma-Aldrich) to quantify biofilm formation at OD_550_ using a spectrophotometer (Tecan Infinite Microplate reader M200 Pro). Motility was assayed by point-inoculating bacterial cultures (OD_600_=0.1) on 0.5x TSB with 0.8% agar and colony sizes were measured after 2 to 3 days. Secretion activities were profiled via point-inoculating (1 µL culture, OD_600_=0.1) bacterial strains on 0.5x TSB agar with 0.1% substrate-of-interest (carbohydrates: pectin, carboxymethyl-cellulose, ⍺-cellulose, xylan, starch; protein: milk and gelatin; lipid: Tween 20), incubated at 28 °C for 2 days. For gelatin, halo of degradation was visualized by incubating the plates in saturated ammonium persulfate for 15 min. For carbohydrates, clearance zones were visualized by staining the plates with 0.1% Congo red (C-6767, SigmaAldrich) for 15 min followed by washing with 6 ppm NaCl solution (0601.1, Roth). All plates were photographed before and after the staining procedures. The enzymatic indices were calculated by dividing the zones of clearing by the colony size

### *In planta* bacterial RNA-Seq

The *in planta Xanthomonas* L148 RNA-Seq was done in accordance to Nobori et al, 2018^61^. Briefly, two-week-old plants grown in agar plates were flood-inoculated with *Xanthomonas* L148 (OD_600_=0.005 in 10 mM MgCl_2_) and shoots of approximately 150 plants were harvested and pooled per sample at 2 dpi when bacterial populations were similar between Col-0 and *rbohD* plants. Samples were harvested, snap-frozen in liquid N_2_, and stored at −80 °C until RNA extraction. The whole experiment was repeated three times. Samples were crushed with metal beads and incubated for 24 h at 4 °C with the isolation buffer^61^. Bacterial cells were separated from the plant tissue via centrifugation. The RNA was isolated from the bacterial pellets using TRIzol (15596026, Invitrogen) and were treated with Turbo DNase (AM1907, Invitrogen) prior to sending to the Max Planck-Genome-Centre Cologne for RNA Sequencing with plant ribo-depletion and cDNA library construction (Universal Prokaryotic RNA-Seq Library Preparation Kit, Tecan) using the Illumina HiSeq 3000 system with 150 bp strand-specific single-end reads resulting in approximately 10 million reads per sample. The resulting reads were mapped to the *Xanthomonas* L148 genome^25^ using the align() function with the default parameters in Rsubread package^62^ to generate BAM files. Mapping rates ranged from 20–46%, which is within the expected values^31^. Mapped reads were counted using DESeq2^63^ using the function featureCounts() from the BAM files and were normalized using the voom() function in limma package^64^ prior to analysis. RNA-Seq raw reads and processed data were deposited in the NCBI GEO repository with accession number GSE226583.

Upon passing quality checks (assessing batch effects through PCA and MA plots for data dispersion), differentially expressed genes were determined using a linear model (gene expression ∼ 0 + genotype + rep; contrast = Col-0 - *rbohD*) and Empirical Bayes statistics with eBayes() function in limma^64^. False discovery rates were accounted for p-values using qvalue^65^. The threshold for significantly differentially expressed genes was set to q-value < 0.05. Principal component analysis was done using the prcomp function^66^; the optimal number of clusters was determined using NbClust() function in NbClust package^67^, cluster memberships were computed with the k-means algorithm^68^, heatmaps were generated using Heatmap() function in ComplexHeatmap package^69^, and pathway enrichment analysis was done for each of the identified gene clusters using enricher() function in clusterProfiler package in R^70^.

### Synthetic community experiment

Two-week-old plants grown in agar plates in a chamber at 23 °C/23°C (day/night) with 10 h of light were flood-inoculated with *Xanthomonas* L148 with or without the leaf-derived synthetic communities (LeafSC, 9 leaf prevalent and functional leaf isolates^27–29^) in two different doses: L148_P1_ + LeafSC contains equal portions of each strain including L148 in the inoculum (Xanthomonas L148/LeafSC, 1:9, each strain would have a final OD_600_=0.01 totaling to OD_600_=0.09 for LeafSC) and L148_P9_ + Leaf SC contains a population of *Xanthomonas* L148 that equals the entire bacterial load of the LeafSC (Xanthomonas L148/LeafSC, 9:9, L148 and the LeafSC at OD_600_=0.09), and were incubated in the same growth chamber. Plants were phenotyped for shoot fresh weights at 14 dpi (See Supplementary Table S2 for list of bacterial strains).

### Protective function experiment

Two-week-old plants grown in agar plates in a chamber at 23 °C/23 °C (day/night) with 10 h of light were flood-inoculated with *Xanthomonas* L148 strains (OD_600_=0.005) and incubated for 5 days. *Pto* lux (OD=0.005) or water was aseptically spray-inoculated (approximately 200 µL per plate) onto the pre-colonized plants. Samples were collected at 0 and 3 dpi to count L148 and *Pto* colonies for different leaf compartments. For the total compartment, leaves were directly homogenized, serially diluted, and plated; for the endophytic compartment, leaves were surface-sterilized with 70% ethanol for 1 min, washed twice with sterile water, homogenized, serially diluted, and then plated. Colonies were allowed to grow on 0.5x TSB agar at 28 °C, and photographs were taken for 1 to 3 days. Colonies were differentiated via their color and chemiluminescence and colonization was expressed as cfu mg^-1^ leaf sample.

### qPCR analysis

Bacterial RNA was isolated from plant samples inoculated with *Xanthomonas* L148 2 dpi or from bacterial pellets from *Xanthomonas* L148 grown in 0.5x TSB with or without 10 µM PQ using TRIzol (15596026, Invitrogen) followed by treatment with Turbo DNase (AM1907, Invitrogen). The cDNA libraries were synthesized with 1 μg RNA input using SuperScript II reverse transcriptase (18064-014, Invitrogen) and random hexamers as primers following the manufacturer’s instructions. An input of 50 ng of cDNA was used for qPCR analyses (CFX Connect Real-Time System, Biorad) of the bacterial genes (please see Supplementary Table S3 for the list of primers and genes tested). The *Δ*Cq was computed by subtracting the Cq of the gene-of-interest from the Cq of the *gyrA* gene from *Xanthomonas* L148.

### Statistical analysis

The R programming environment (R version 4.2.2) was used for data analysis and visualization^66^. The data were inspected for the assumptions of the linear model (homodescacity, independence, and normality) and were normalized, if necessary, prior to statistical analysis using ANOVA with *post hoc* Tukey’s HSD test or the Least Significant Difference (LSD) test using the package agricolae^71^.

### Genomic interrogation for CAZyme functions

Genomes for *Xanthomonas* L148 and other Xanthomonadales strains within the AtSPHERE^25^ and known *Xanthomonas* pathogens (downloaded from NCBI; Sayers, et al, 2022) were annotated for CAZyme functions (http://www.cazy.org/)^72^ using the eggnog mapper (http://eggnog-mapper.embl.de/)^73^ to determine the CAZyme repertoire of the bacterial strains and their potential substrates.

### Data deposition

The *in planta* bacterial RNA-Seq data reported in this paper have been deposited in the Gene Expression Omnibus (GEO) database, https://www.ncbi.nlm.nih.gov/geo (accession no. GSE226583).

## Code availability

No custom code was generated for this study.

## Acknowledgements

We thank Neysan Donnelly for editing and Wanqing Jiang for providing helpful comments on the manuscript. This work was supported by the National Key R&D Program of China (2022YFA1304403 to K.T.), the National Natural Science Foundation of China (32250710139 to K.T.), Joint Funding of Huazhong Agricultural University and Agricultural Genomics Institute at Shenzhen, Chinese Academy of Agricultural Sciences (SZYJY2021007 to K.T. and X.H.), the Max Planck Society (to P.S.-L and K.T.), and a German Research Foundation (DFG) grant (SPP2125) (to P.S.-L and K.T.).

## Author Contribution

F.E. and K.T. conceived the research. F.E., X.H., P.S.-L, and K.T. designed the research. A.M. designed and constructed *Pto* lux. F.E. performed all of the experimental work and the analysis of the data. F.E. and K.T. wrote the manuscript with input from all the authors.

## List of Supplementary Tables and Datasets

**Supplementary Table S1.** List of *Arabidopsis thaliana* wild-type and mutants used in this study

**Supplementary Table S2.** List of bacterial strains used and generated in this study.

**Supplementary Table S3.** List of primers and PCR profiles used in this study

**Supplementary Dataset S1.** List of *Xanthomonas* L148::Tn5 mutant candidates with loss-of-mortality in *rbohD* phenotypes using the high-throughput screening.

**Supplementary Dataset S2.** Top table of the DEGs for *in planta Xanthomonas* L148 transcriptome Col-0 vs. *rbohD* colonized plants.

**Supplementary Dataset S3.** Clustering membership of the DEGs and the GO term enrichment analysis for the gene clusters.

**Supplementary Figure S1.**
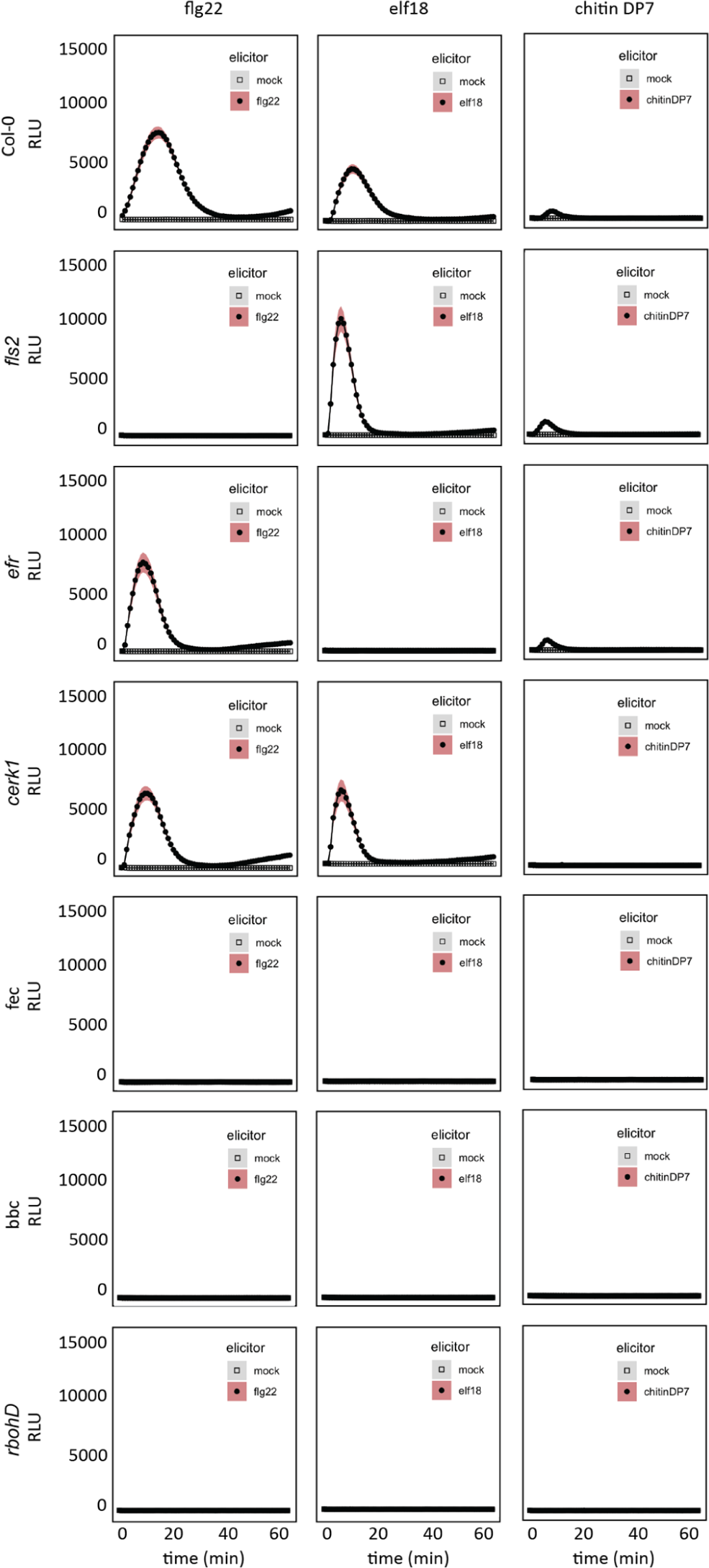
ROS burst profile of immune-compromised mutants and Col-0 wildtype plants with MAMPs. Leaf discs from 5-to-6-week-old plants were treated with 1 µM of MAMPs, flg22, elf18, and chitinDP7. The immune-compromised mutant *fls2* lacks the receptor recognizing flg22, *efr* lacks the receptor for elf18, and *cerk1* lacks the co-receptor for chitinDP7; *fec* (*fls2 efr, cerk1*) and *bbc* (*bak1 bkk1 cerk1*) are triple mutants lacking the MAMP (co) receptor. Data from at least 2 independent experiments each with 8 biological replicates were used.

**Supplementary Figure S2.**
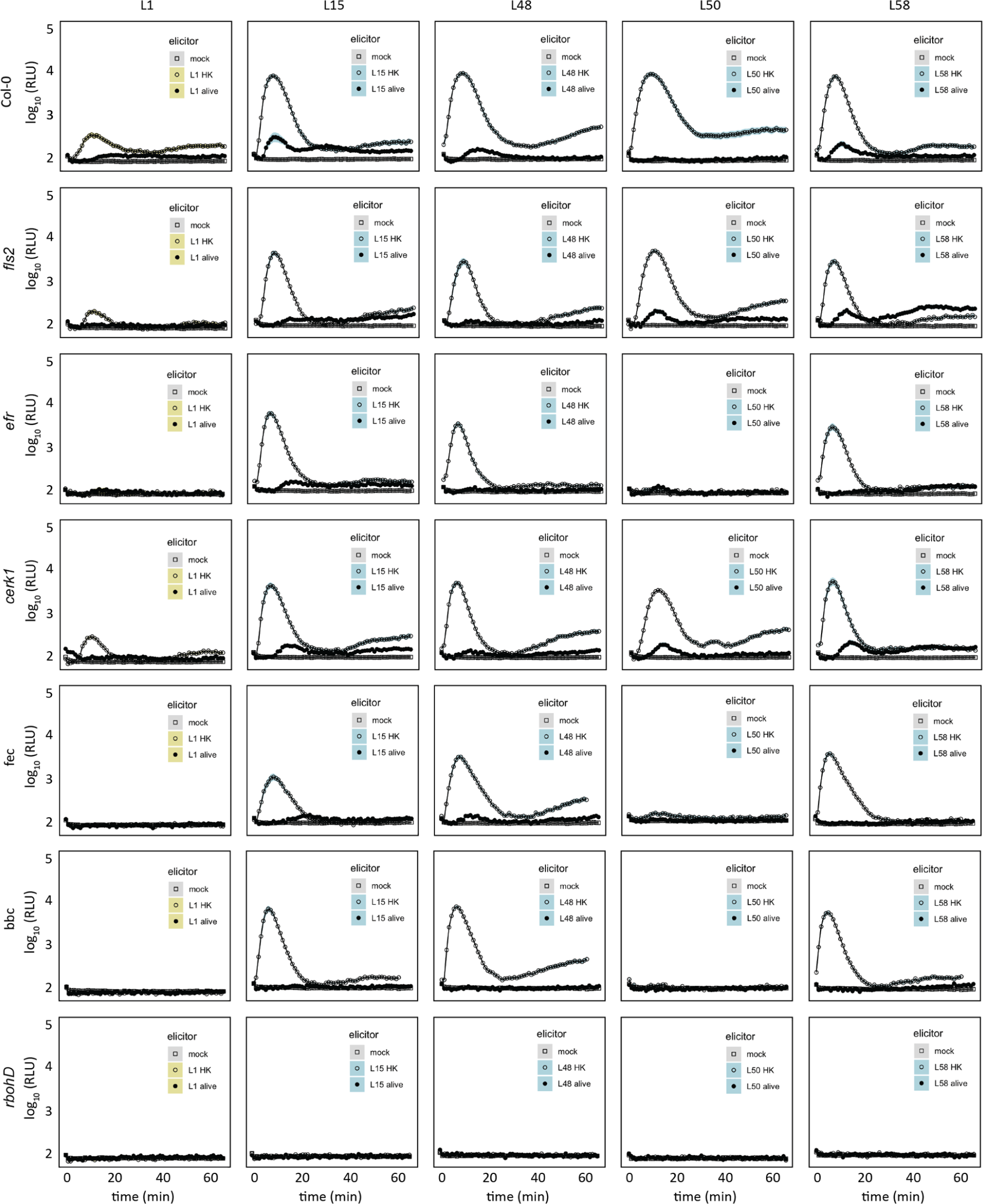
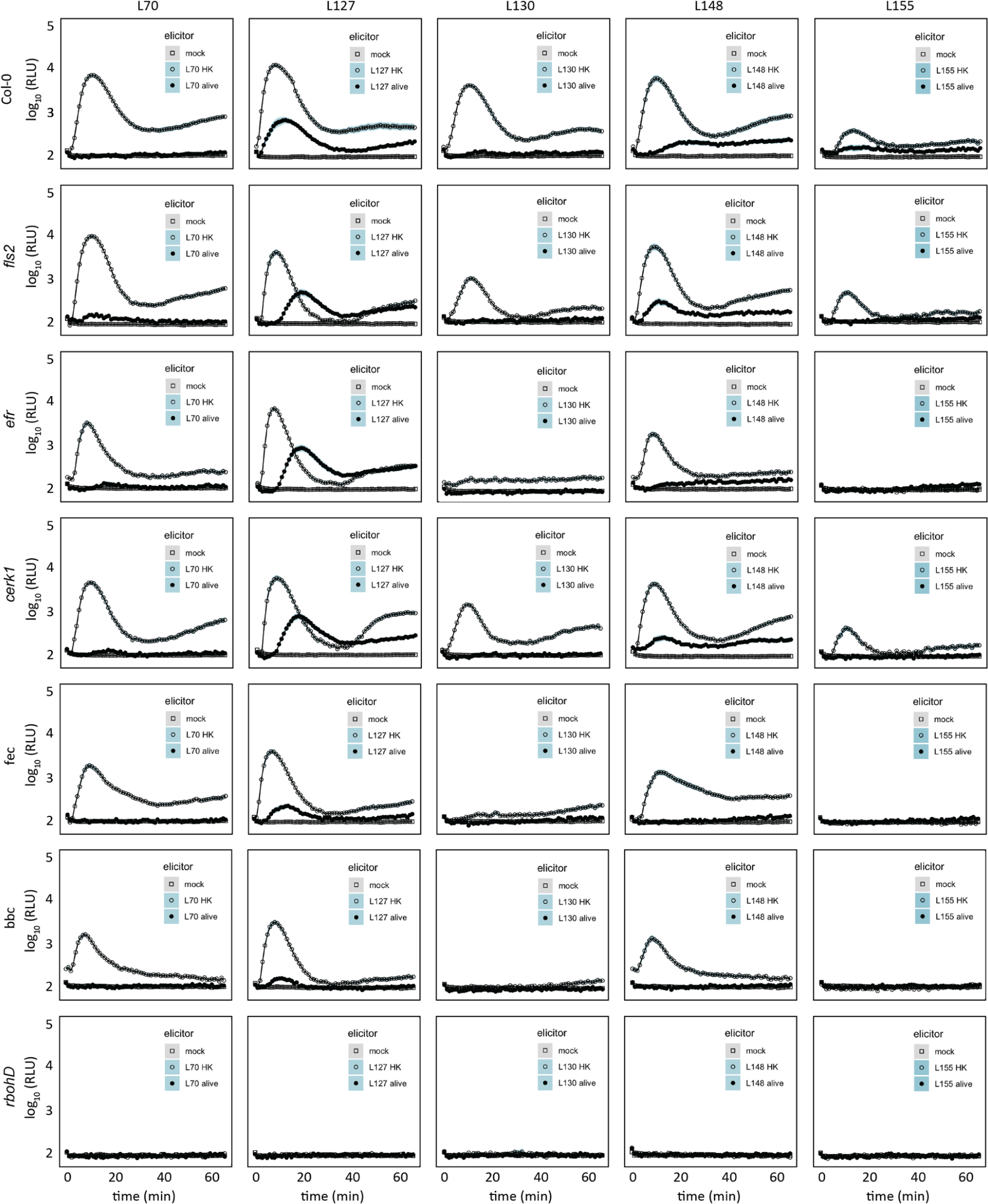
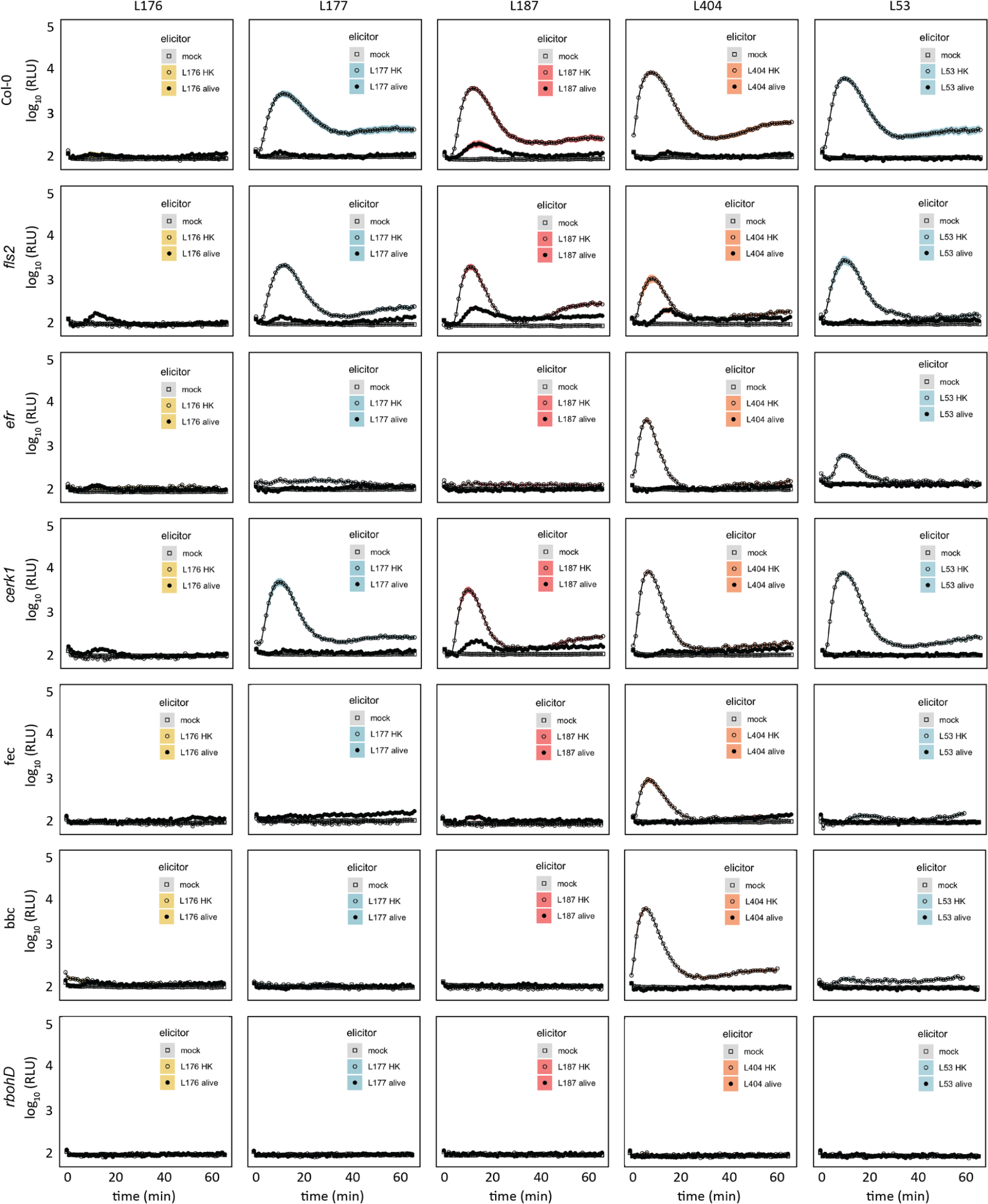
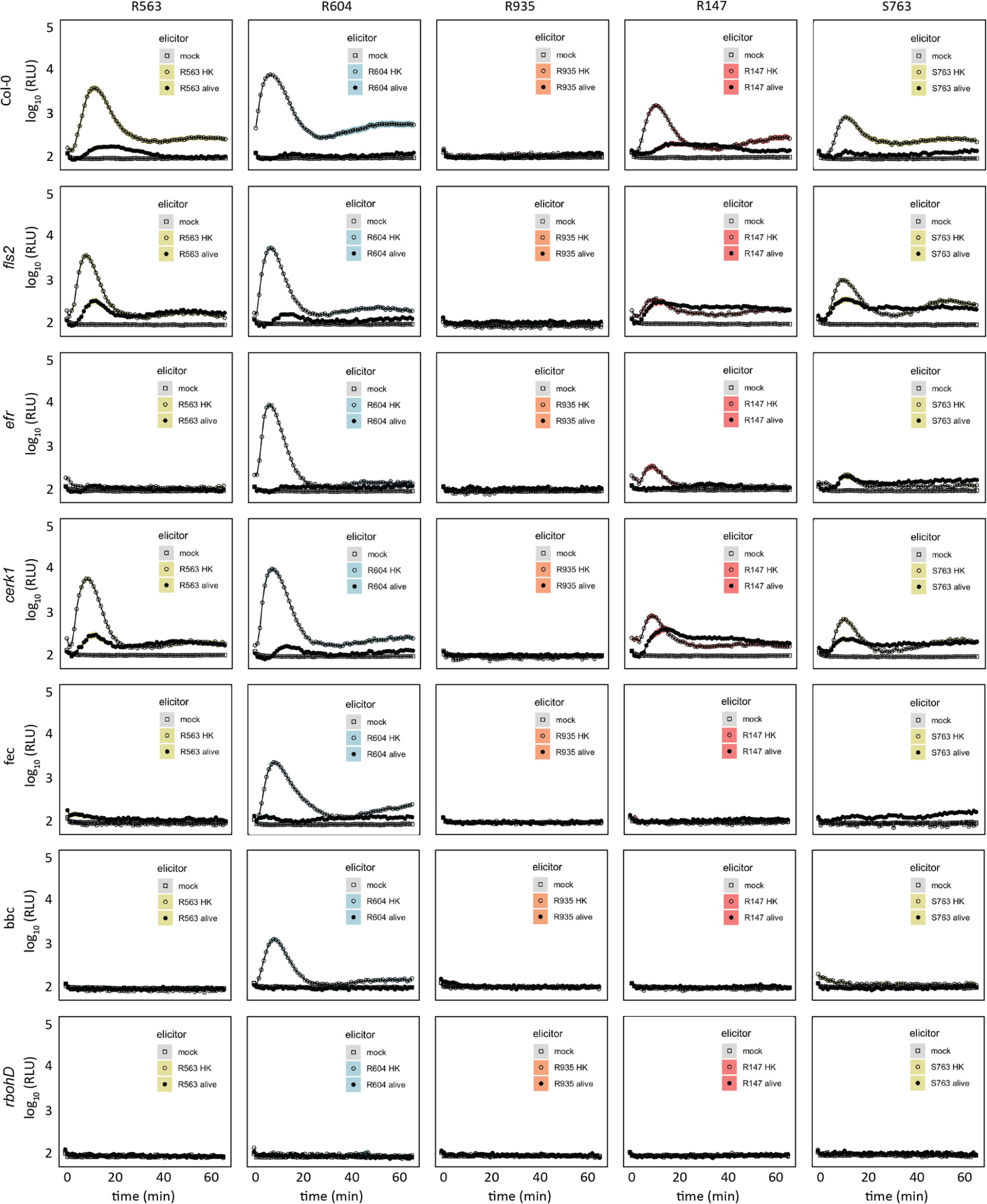
ROS burst profile of immune-compromised mutants and Col-0 wild-type plants with commensal bacteria. Leaf discs from 5-to-6-week-old plants were inoculated with live or heat-killed microbiota strains (OD600=0.5) in mono-associations for ROS burst assays. Data from at least 2 independent experiments each with 8 biological replicates were used.

**Supplementary Figure S3.**
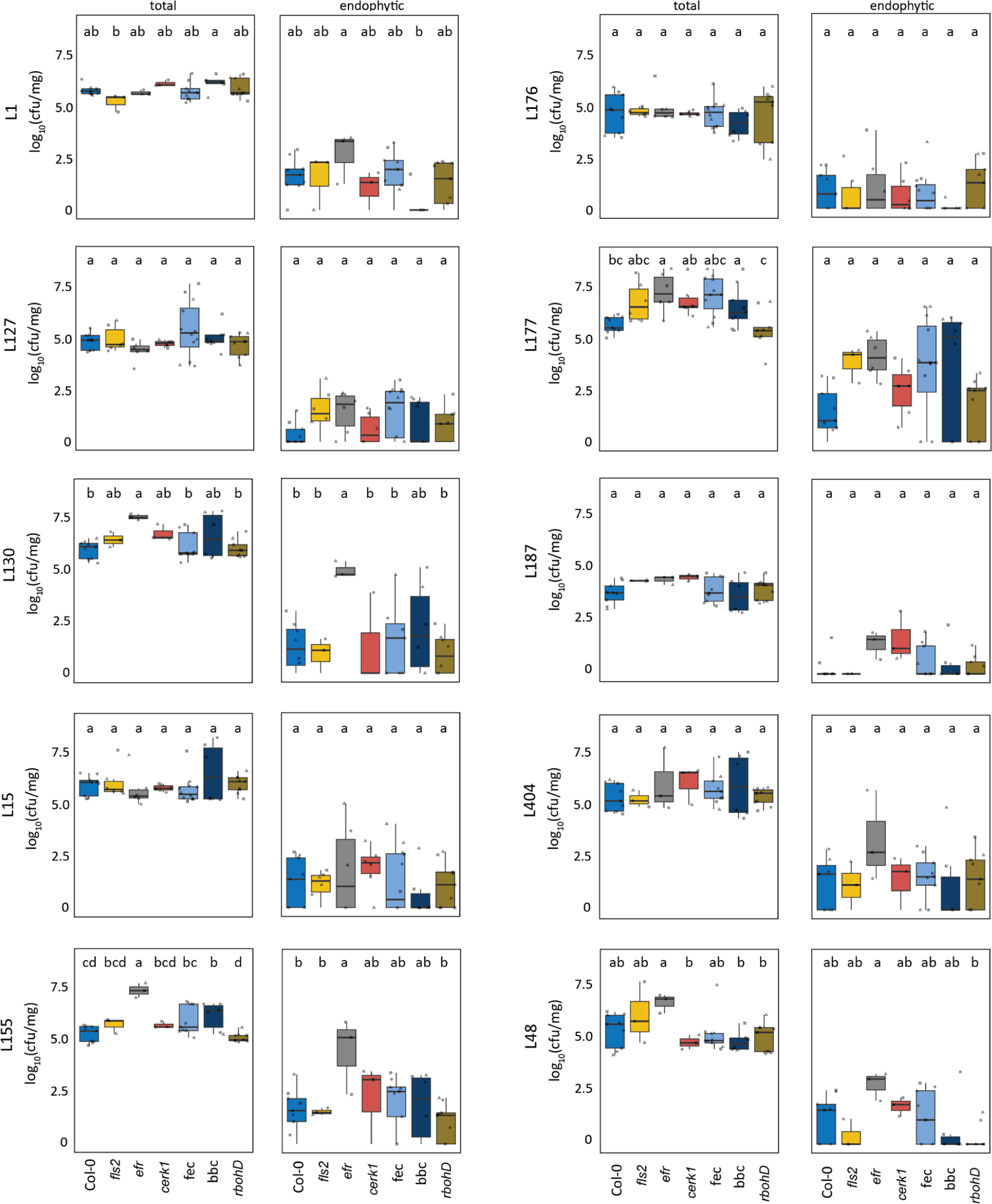
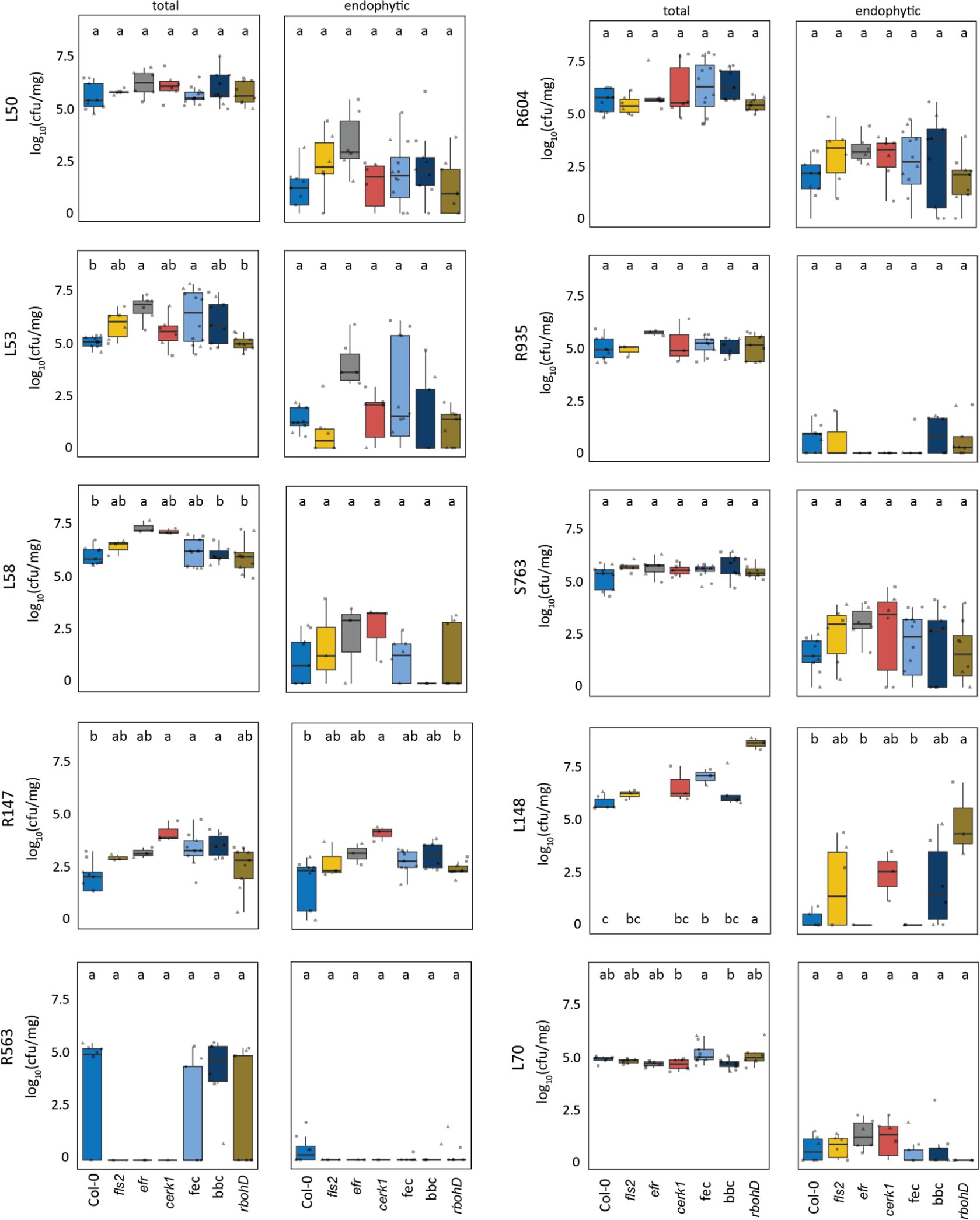
Leaf colonization capacities of commensal bacteria on immune-compromised mutants and Col-0 wildtype plants. Two-week-old axenic plants were flood-inoculated with microbiota strains (OD600=0.005) and were plated for colony counts for the total and endophytic leaf compartments at 5 dpi. Data from at least 2 independent experiments each with 8 biological replicates were used. Different letters indicate statistically significant differences (ANOVA with *post hoc* Tukey’s test, *P* ≤ 0.05). Results are depicted as box plots with the boxes spanning the interquartile range (IQR, 25^th^ to 75^th^ percentiles), the mid-line indicates the median, and the whiskers cover the minimum and maximum values not extending beyond 1.5x of the IQR.

**Supplementary Figure S4.**
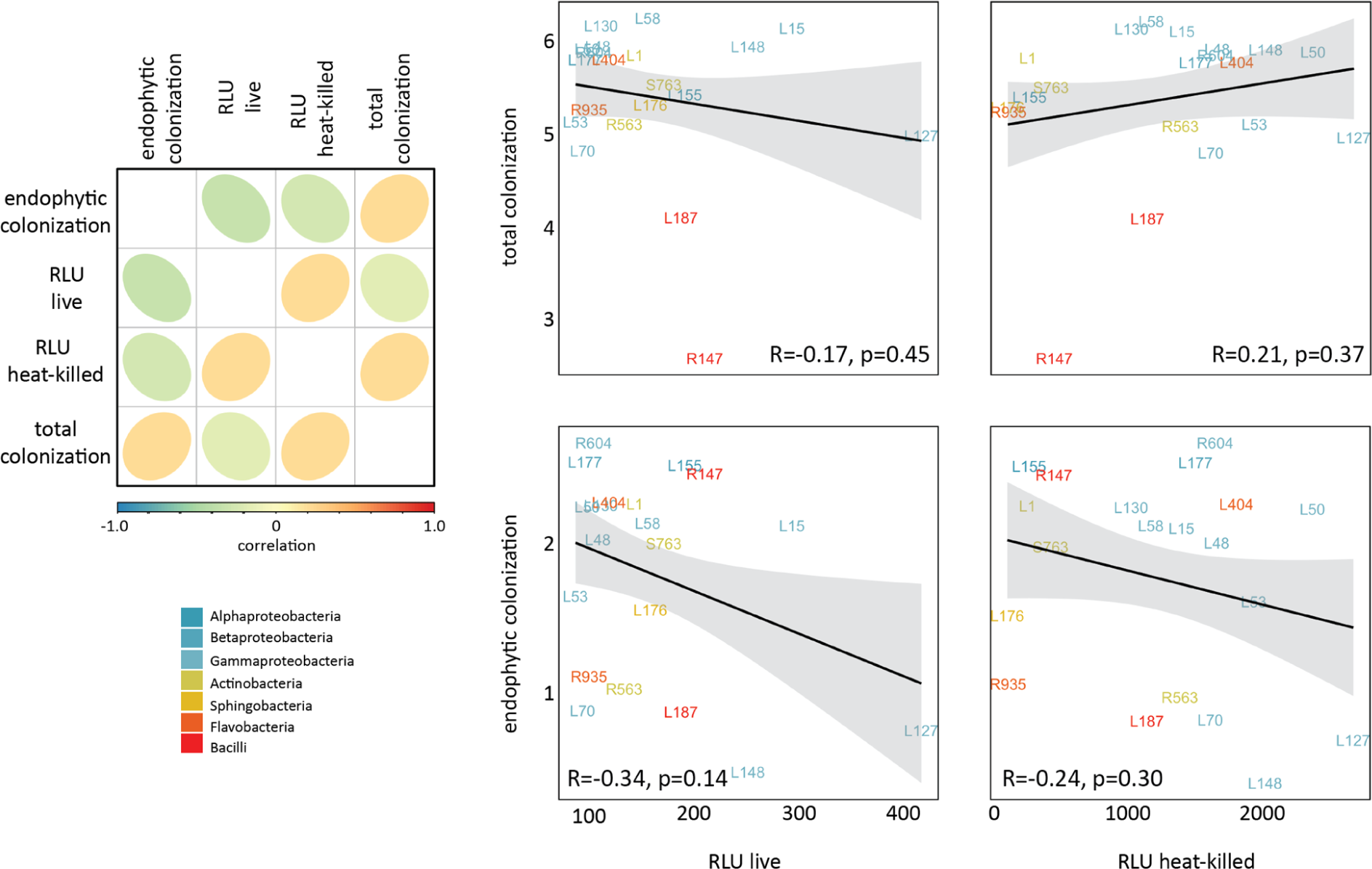
The ROS outburst profile and colonization capacities in Col-0 wild-type plants with the microbiota members have poor correlation. Correlational analysis of the capacity of the strain (live or heat-killed versions) to induce ROS and their corresponding colonization profiles in wild-type Col-0 plants (R, coefficient of determination, p ≤ 0.02). For ROS outburst assay, leaf discs from 5-to-6-week-old plants were triggered with live or heat-killed microbiota strains (OD600=0.5) in mono-associations. For colonization assays, two-week-old axenic plants were flood-inoculated with microbiota strains (OD600=0.005) and were plated for colony counts for the total and endophytic leaf compartments at 5 dpi.

**Supplementary Figure S5.**
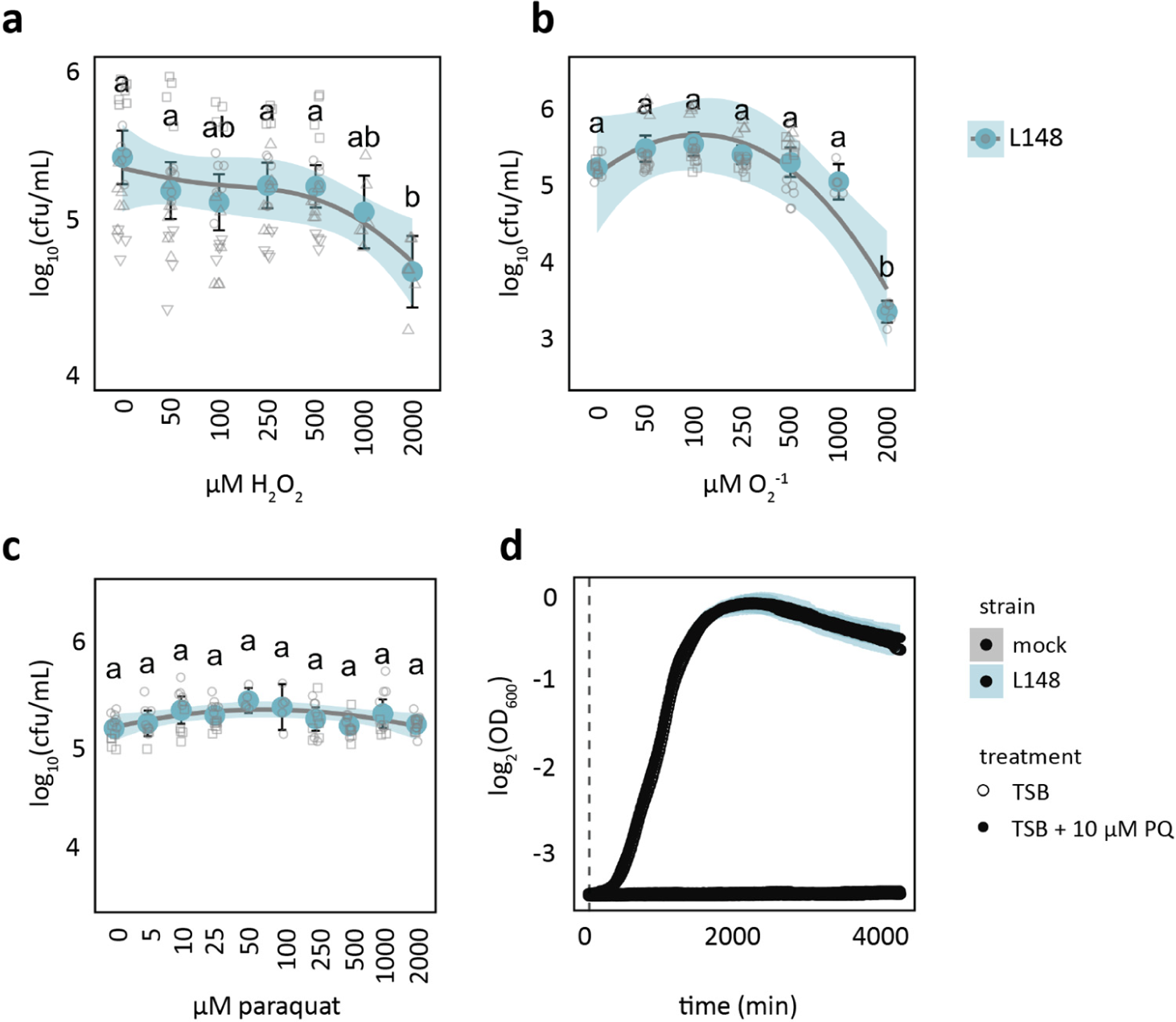
*Xanthomonas* L148 is not sensitive to *in vitro* exposure to ROS compounds. **a–c**, Recovery of *Xanthomonas* L148 bacterial cells (initial inoculum OD600 = 0.02) upon acute exposure with ROS compounds H2O2 (**a**), O2^-^ (**b**), and PQ (**c**) in different concentrations (0–2000 µM). H2O2 was applied at different doses for 30 min. For O2^-^ treatment, 1 mol of xanthine is converted to 1 mol O2^-1^ with 1 U xanthine oxidase at pH 7.5 at 25 °C in 1 min; reactions were commenced and bacterial cells were sampled at different time points: 0, 2, 4, 10, 20, 60, and 80 min to produce 0, 50, 100, 250, 500, 1000, and 2000 μM O2^-^, respectively. **d**, Growth curves of *Xanthomonas* L148 in TSB upon chronic exposure of 0 or 10 µM PQ for 4000 min. **a-d**, Data were from at least 2 independent experiments each with 3–4 biological replicates. Different letters indicate statistically significant differences (ANOVA with *post hoc* Tukey’s test, *P* ≤ 0.05).

**Supplementary Figure S6.**
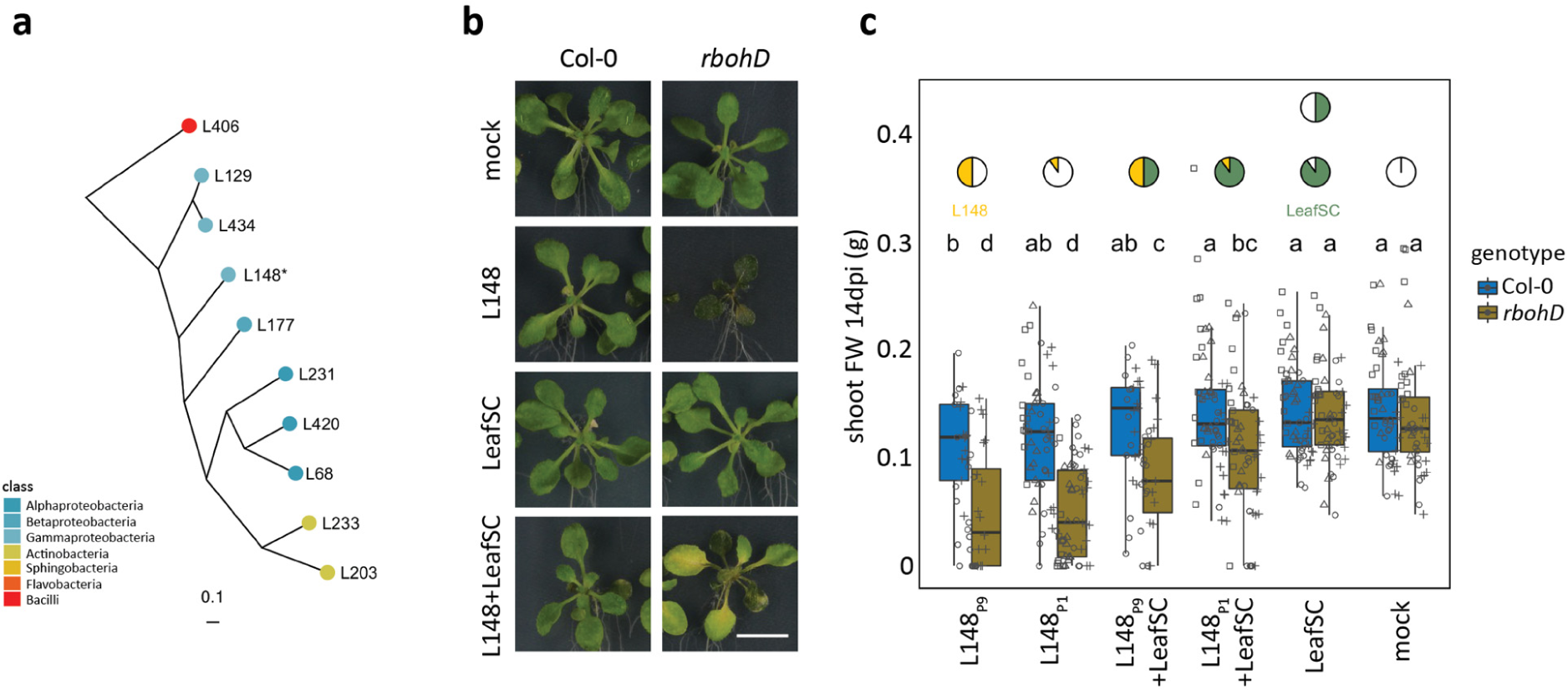
*Xanthomonas* L148 pathogenic potential was partially suppressed by the presence of other leaf commensals. **a**, Phylogenetic relationship of the strains comprising the leaf-derived synthetic community (LeafSC) which consists of strains that are robust and prevalent leaf colonizers, and taxonomically represents diverse members of the leaf microbiota. **b, c**, Representative image (**b**) and the measured shoot fresh weights (**c**) of Col-0 and *rbohD* plants flood-inoculated with mock, LeafSC, L148P1 + LeafSC (equal portions of *Xanthomonas* L148 with each strain: L148/LeafSC, 1:9, final OD600=0.01), L148P9 + LeafSC (portion of *Xanthomonas* L148 equals the bacterial load of the all strains: L148/LeafSC, 9:9, final OD600=0.01), and the equivalent doses of *Xanthomonas* L148 (L148P1 and L148P9, P9 is 9 times the dose of P1). The pies indicate the relative proportion of the *Xanthomonas* L148 = yellow and LeafSC = green. White horizontal bar = 1 cm. Data from 2 independent experiments each with 3–4 replicates were used. Different letters indicate statistically significant differences (ANOVA with *post hoc* Tukey’s test, *P* ≤ 0.05).

**Supplementary Figure S7.**
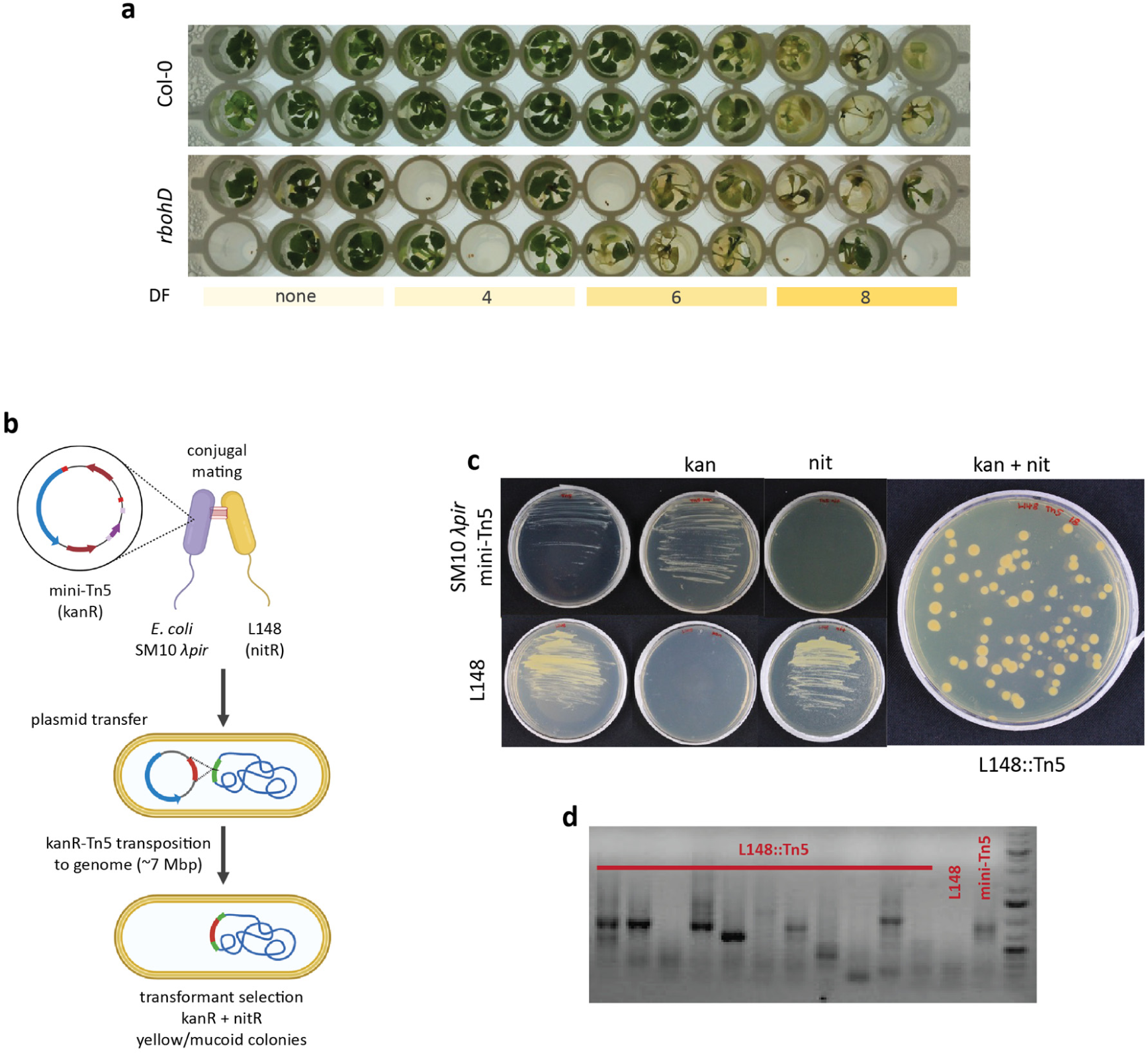
Optimization of high-throughput genome-wide screening and generation of the *Xanthomonas* L148::Tn5 mutant library. **a**, Representative image of Col-0 wild-type and *rbohD* mutant plants inoculated with serially diluted *Xanthomonas* L148 suspensions in the high-throughput 96-well plate format. A dilution factor (DF) of 6 was chosen for the best contrast between Col-0 and *rbohD*. **b**, Schematic diagram of the construction of the *Xanthomonas* L148::Tn5 mutant library via conjugation with *E. coli* harboring the mini-Tn5 plasmid. **c**, Antibiotic resistance of *Xanthomonas* L148, *E. coli* SM10λpir and the *Xanthomonas* L148::Tn5 mutants. The parental strain *Xanthomonas* L148 is resistant to nitrofurantoin (nit, 50 μg/mL in TSB medium) which was used for counter-selection for the plasmid carrier *E. coli.* The mini-Tn5 carrying *E. coli* is resistant to kanamycin (kan, 50 μg/mL in TSB medium) and was used for selecting against the wild-type *Xanthomonas* L148. *Xanthomonas* L148::Tn5 transformants are resistant to both nit and kan in TSB medium. **d**, Electrophoretogram of the genomic transposon insertion PCR validation for the randomly selected *Xanthomonas* L148::Tn5 mutant strains. PCR products were Sanger-sequenced to determine the transposon insertion site.

**Supplementary Figure S8.**
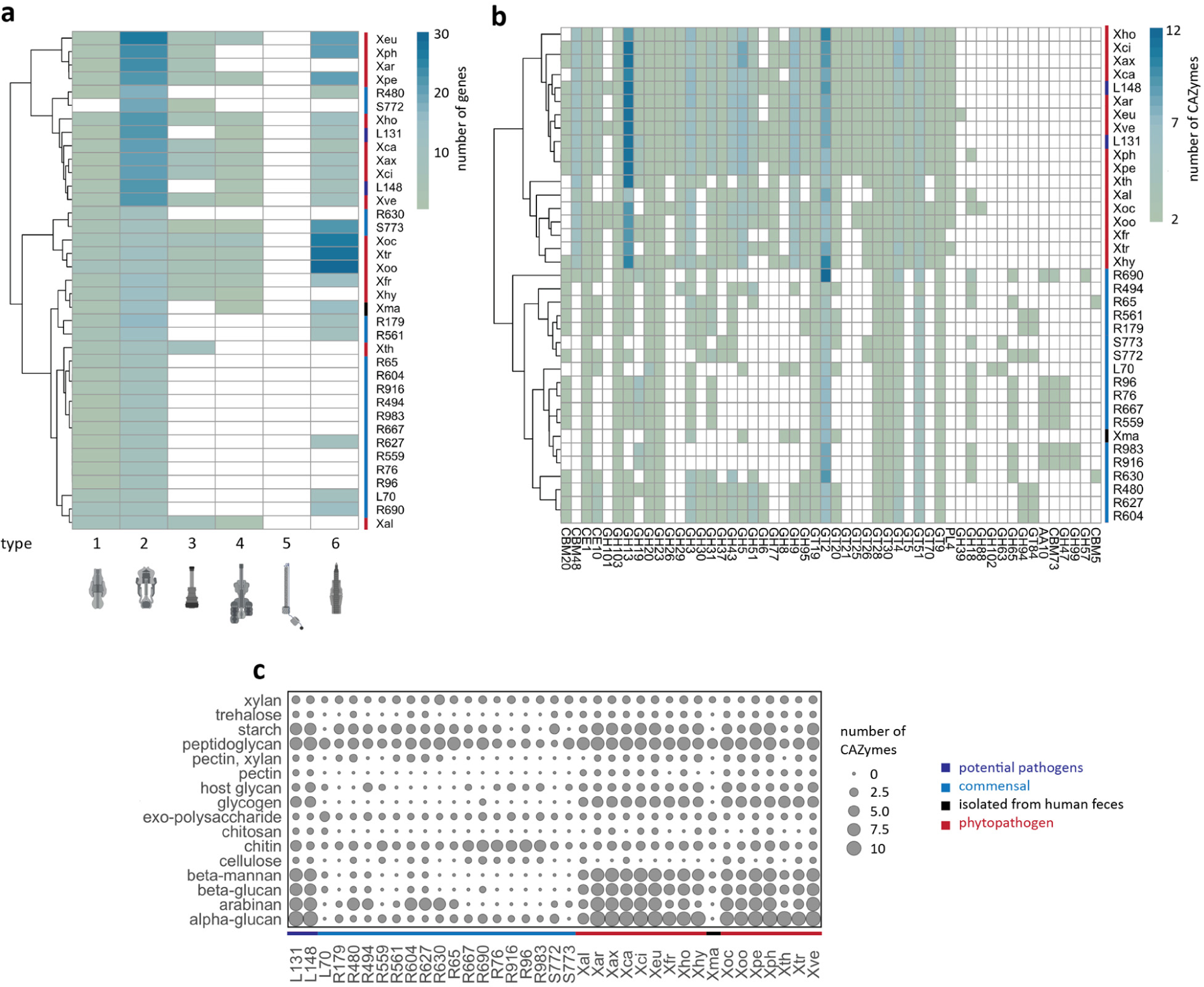
Secretion systems and CAZyme repertoire of Xanthomonadales clade. **a-b**, genomic examination of Xanthomonadales members of *A. thaliana* microbiota (20) and pathogenic *Xanthomonas* strains (17): Xal, = *X. albineans*; Xar = *X. arboricola;* Xax = *X. axonopodis*; Xca = *X. campestris;* Xci = *X. citri;* Xeu = *X. euvesicatoria;* Xfr = *X. fragariae;* Xho = *X. hortorum;* Xhy = *X. hyacinthi;* Xoc = *X. oryzae* pv. *oryzicola*; Xoo = *X. oryzae* pv. *oryzae;* Xpe = *X. perforans;* Xph = *X. phaseoli;* Xth = *X. theicola;* Xtr = *X. translucens;* Xve = *X. vesicatoria.* Xma = *X. massiliensis* is non-pathogenic strain isolated from human feces; L148 (in this study) and L131 (Pfeilmeier et al, 2021) are potentially pathogenic. **a**, occurrence of type 1 to 6 secretion systems. **b**, CAZyme repertoire of the Xanthomonadales. **c**, potential substrates of the genome encoded CAZymes. Some illustrations created in BioRender.

**Supplementary Figure S9.**
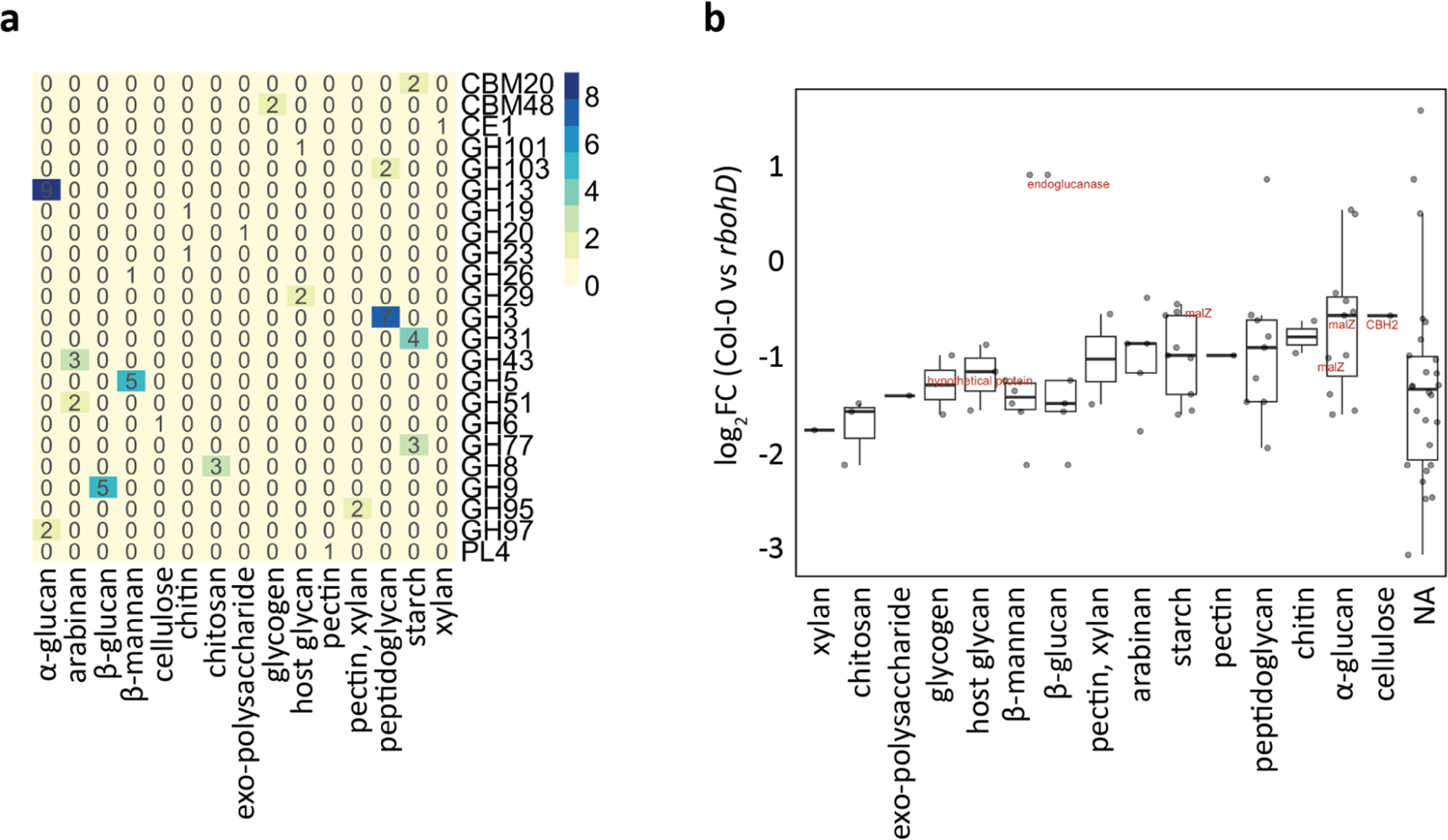
Significantly down-regulated CAZymes in Col-0 can potentially degrade plant host cell wall components. **a**, Heatmap representation of the number of significantly differentially expressed CAZyme genes in *Xanthomonas* L148 (Col-0 vs. *rbohD*) with the respective potential substrates. **b**, Log2 fold changes of the CAZyme gene expression (Col-0 vs. *rbohD*) with their respective potential substrates; *Xanthomonas* L148::Tn5 candidate genes with CAZyme annotations are labelled. Results in **b** are depicted as box plots with the boxes spanning the interquartile range (IQR, 25^th^ to 75^th^ percentiles), the mid-line indicates the median, and the whiskers cover the minimum and maximum values not extending beyond 1.5x of the IQR.

**Supplementary Figure S10.**
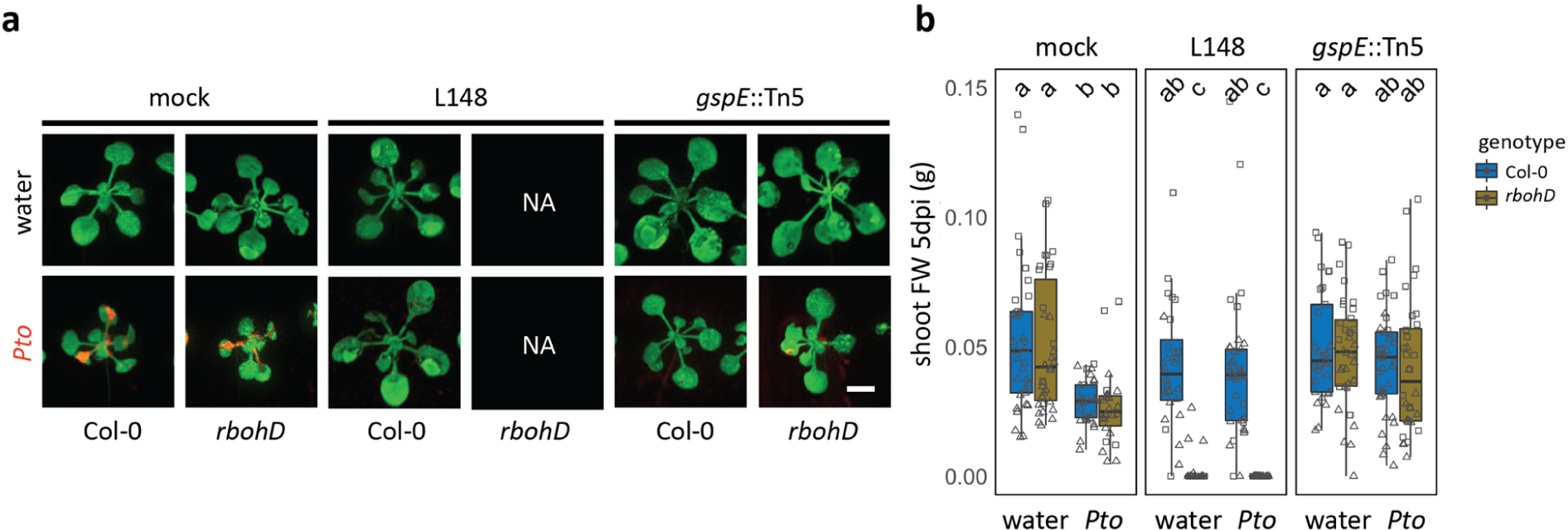
Performance of plants pre-colonized with commensals after pathogen invasion. **a,b**, Representative images (**a**) and quantification of shoot fresh weight as a plant health parameter (**b**). 14-day-old Col-0 and *rbohD* plants grown on agar plates were flood-inoculated with wildtype *Xanthomonas* L148 and *gspE*::Tn5 (OD600=0.005) for 5 days followed by spray infection with *Pto*. Samples were taken at 5 dpi (2 independent experiments each with 3–5 biological replicates). Red patches in the images indicates colonization by the pathogen. Different letters indicate statistically significant differences (ANOVA with *post hoc* Tukey’s test, *P* ≤ 0.05). Results in **b** are depicted as box plots with the boxes spanning the interquartile range (IQR, 25^th^ to 75^th^ percentiles), the mid-line indicates the median, and the whiskers cover the minimum and maximum values not extending beyond 1.5x of the IQR.

## References

1 Müller, D. B., Vogel, C., Bai, Y., & Vorholt, J. A. (2016). The Plant Microbiota: Systems-Level Insights and Perspectives. Annual review of genetics, 50, 211–234. https://doi.org/10.1146/annurev-genet-120215-034952

2 Drew, G. C., Stevens, E. J., & King, K. C. (2021). Microbial evolution and transitions along the parasite-mutualist continuum. Nature reviews. Microbiology, 19(10), 623–638. https://doi.org/10.1038/s41579-021-00550-7

3 Jochum, L., & Stecher, B. (2020). Label or Concept - What Is a Pathobiont?. Trends in microbiology, 28(10), 789–792. https://doi.org/10.1016/j.tim.2020.04.011

4 Caballero-Flores, G., Pickard, J. M., & Núñez, G. (2022). Microbiota-mediated colonization resistance: mechanisms and regulation. Nature reviews. Microbiology, 10.1038/s41579-022-00833-7. Advance online publication. https://doi.org/10.1038/s41579-022-00833-7

5 Dodds, P. N., & Rathjen, J. P. (2010). Plant immunity: towards an integrated view of plant-pathogen interactions. Nature reviews. Genetics, 11(8), 539–548. https://doi.org/10.1038/nrg2812

6 Zipfel, C., Robatzek, S., Navarro, L., Oakeley, E. J., Jones, J. D., Felix, G., & Boller, T. (2004). Bacterial disease resistance in Arabidopsis through flagellin perception. Nature, 428(6984), 764–767. https://doi.org/10.1038/nature02485

7 Zipfel, C., Kunze, G., Chinchilla, D., Caniard, A., Jones, J. D., Boller, T., & Felix, G. (2006). Perception of the bacterial PAMP EF-Tu by the receptor EFR restricts Agrobacterium-mediated transformation. Cell, 125(4), 749–760. https://doi.org/10.1016/j.cell.2006.03.037

8 Roux, M., Schwessinger, B., Albrecht, C., Chinchilla, D., Jones, A., Holton, N., Malinovsky, F. G., Tör, M., de Vries, S., & Zipfel, C. (2011). The Arabidopsis leucine-rich repeat receptor-like kinases BAK1/SERK3 and BKK1/SERK4 are required for innate immunity to hemibiotrophic and biotrophic pathogens. The Plant cell, 23(6), 2440–2455. https://doi.org/10.1105/tpc.111.084301

9 Miya, A., Albert, P., Shinya, T., Desaki, Y., Ichimura, K., Shirasu, K., Narusaka, Y., Kawakami, N., Kaku, H., & Shibuya, N. (2007). CERK1, a LysM receptor kinase, is essential for chitin elicitor signaling in Arabidopsis. Proceedings of the National Academy of Sciences of the United States of America, 104(49), 19613–19618. https://doi.org/10.1073/pnas.0705147104

10 Boller, T., & Felix, G. (2009). A renaissance of elicitors: perception of microbe-associated molecular patterns and danger signals by pattern-recognition receptors. Annual review of plant biology, 60, 379–406. https://doi.org/10.1146/annurev.arplant.57.032905.105346

11 Xin, X. F., Nomura, K., Aung, K., Velásquez, A. C., Yao, J., Boutrot, F., Chang, J. H., Zipfel, C., & He, S. Y. (2016). Bacteria establish an aqueous living space in plants crucial for virulence. Nature, 539(7630), 524–529. https://doi.org/10.1038/nature20166

12 Chen, T., Nomura, K., Wang, X. et al. A plant genetic network for preventing dysbiosis in the phyllosphere. Nature 580, 653–657 (2020). https://doi.org/10.1038/s41586-020-2185-0

13 Torres, M. A., Dangl, J. L., & Jones, J. D. (2002). Arabidopsis gp91phox homologues AtrbohD and AtrbohF are required for accumulation of reactive oxygen intermediates in the plant defense response. Proceedings of the National Academy of Sciences of the United States of America, 99(1), 517–522. https://doi.org/10.1073/pnas.012452499

14 Castro, B., Citterico, M., Kimura, S., Stevens, D. M., Wrzaczek, M., & Coaker, G. (2021). Stress-induced reactive oxygen species compartmentalization, perception and signalling. Nature plants, 7(4), 403–412. https://doi.org/10.1038/s41477-021-00887-0

15 Bolwell, G.P., Daudi, A. (2009). Reactive Oxygen Species in Plant–Pathogen Interactions. In: Rio, L., Puppo, A. (eds) Reactive Oxygen Species in Plant Signaling. Signaling and Communication in Plants. Springer, Berlin, Heidelberg. https://doi.org/10.1007/978-3-642-00390-5_7

16 Song, Y., Wilson, A. J., Zhang, X. C., Thoms, D., Sohrabi, R., Song, S., Geissmann, Q., Liu, Y., Walgren, L., He, S. Y., & Haney, C. H. (2021). FERONIA restricts Pseudomonas in the rhizosphere microbiome via regulation of reactive oxygen species. Nature plants, 7(5), 644– 654. https://doi.org/10.1038/s41477-021-00914-0

17 Pfeilmeier, S., Petti, G. C., Bortfeld-Miller, M., Daniel, B., Field, C. M., Sunagawa, S., & Vorholt, J. A. (2021). The plant NADPH oxidase RBOHD is required for microbiota homeostasis in leaves. Nature microbiology, 6(7), 852–864. https://doi.org/10.1038/s41564-021-00929-5

18 Tzipilevich, E., Russ, D., Dangl, J. L., & Benfey, P. N. (2021). Plant immune system activation is necessary for efficient root colonization by auxin-secreting beneficial bacteria. Cell host & microbe, 29(10), 1507–1520.e4. https://doi.org/10.1016/j.chom.2021.09.005

19 Kadota, Y., Sklenar, J., Derbyshire, P., Stransfeld, L., Asai, S., Ntoukakis, V., Jones, J. D., Shirasu, K., Menke, F., Jones, A., & Zipfel, C. (2014). Direct regulation of the NADPH oxidase RBOHD by the PRR-associated kinase BIK1 during plant immunity. Molecular cell, 54(1), 43–55. https://doi.org/10.1016/j.molcel.2014.02.021

20 Salmond, G. P. (1994). Secretion of extracellular virulence factors by plant pathogenic bacteria. Annual review of phytopathology, 32(1), 181–200.

21 Tampakaki A. P. (2014). Commonalities and differences of T3SSs in rhizobia and plant pathogenic bacteria. Frontiers in plant science, 5, 114. https://doi.org/10.3389/fpls.2014.00114

22 Kambara, K., Ardissone, S., Kobayashi, H., Saad, M. M., Schumpp, O., Broughton, W. J., & Deakin, W. J. (2009). Rhizobia utilize pathogen-like effector proteins during symbiosis. Molecular microbiology, 71(1), 92–106. https://doi.org/10.1111/j.1365-2958.2008.06507.x

23 Maekawa, T., Kufer, T. A., & Schulze-Lefert, P. (2011). NLR functions in plant and animal immune systems: so far and yet so close. Nature immunology, 12(9), 817–826. https://doi.org/10.1038/ni.2083

24 Teixeira, P. J. P. L., Colaianni, N. R., Law, T. F., Conway, J. M., Gilbert, S., Li, H., Salas-González, I., Panda, D., Del Risco, N. M., Finkel, O. M., Castrillo, G., Mieczkowski, P., Jones, C. D., & Dangl, J. L. (2021). Specific modulation of the root immune system by a community of commensal bacteria. Proceedings of the National Academy of Sciences of the United States of America, 118(16), e2100678118. https://doi.org/10.1073/pnas.2100678118

25 Bai, Y., Müller, D., Srinivas, G. et al. (2015). Functional overlap of the *Arabidopsis* leaf and root microbiota. Nature 528, 364–369, https://doi.org/10.1038/nature16192

26 Nobori, T., Cao, Y., Entila, F., Dahms, E., Tsuda, Y., Garrido-Oter, R., & Tsuda, K. (2022). Dissecting the cotranscriptome landscape of plants and their microbiota. EMBO reports, 23(12), e55380. https://doi.org/10.15252/embr.202255380

27 Carlström, C.I., Field, C.M., Bortfeld-Miller, M. et al. Synthetic microbiota reveal priority effects and keystone strains in the *Arabidopsis* phyllosphere. Nat Ecol Evol 3, 1445–1454 (2019). https://doi.org/10.1038/s41559-019-0994-z

28 Thiergart, T., Durán, P., Ellis, T. et al. Root microbiota assembly and adaptive differentiation among European *Arabidopsis* populations. Nat Ecol Evol 4, 122–131 (2020). https://doi.org/10.1038/s41559-019-1063-3

29 Karasov, T. L., Almario, J., Friedemann, C., Ding, W., Giolai, M., Heavens, D., Kersten, S., Lundberg, D. S., Neumann, M., Regalado, J., Neher, R. A., Kemen, E., & Weigel, D. (2018). Arabidopsis thaliana and Pseudomonas Pathogens Exhibit Stable Associations over Evolutionary Timescales. Cell host & microbe, 24(1), 168–179.e4. https://doi.org/10.1016/j.chom.2018.06.011

30 Cianciotto N. P. (2005). Type II secretion: a protein secretion system for all seasons. Trends in microbiology, 13(12), 581–588. https://doi.org/10.1016/j.tim.2005.09.005

31 Nobori, T., Velásquez, A. C., Wu, J., Kvitko, B. H., Kremer, J. M., Wang, Y., He, S. Y., & Tsuda, K. (2018). Transcriptome landscape of a bacterial pathogen under plant immunity. Proceedings of the National Academy of Sciences of the United States of America, 115(13), E3055–E3064. https://doi.org/10.1073/pnas.1800529115

32 Vogel, C. M., Potthoff, D. B., Schäfer, M., Barandun, N., & Vorholt, J. A. (2021). Protective role of the Arabidopsis leaf microbiota against a bacterial pathogen. Nature microbiology, 6(12), 1537–1548. https://doi.org/10.1038/s41564-021-00997-7

33 Jakob, K., Goss, E. M., Araki, H., Van, T., Kreitman, M., & Bergelson, J. (2002). Pseudomonas viridiflava and P. syringae--natural pathogens of Arabidopsis thaliana. Molecular plant-microbe interactions: MPMI, 15(12), 1195–1203. https://doi.org/10.1094/MPMI.2002.15.12.1195

34 Agler MT, Ruhe J, Kroll S, Morhenn C, Kim S-T, Weigel D, et al. (2016) Microbial Hub Taxa Link Host and Abiotic Factors to Plant Microbiome Variation. PLoS Biol 14(1): e1002352. https://doi.org/10.1371/journal.pbio.1002352

35 Durán, P., Thiergart, T., Garrido-Oter, R., Agler, M., Kemen, E., Schulze-Lefert, P., & Hacquard, S. (2018). Microbial Interkingdom Interactions in Roots Promote Arabidopsis Survival. Cell, 175(4), 973–983.e14. https://doi.org/10.1016/j.cell.2018.10.020

36 Ma, KW., Niu, Y., Jia, Y. et al. Coordination of microbe–host homeostasis by crosstalk with plant innate immunity. Nat. Plants 7, 814–825 (2021). https://doi.org/10.1038/s41477-021-00920-2

37 Shalev, O., Karasov, T.L., Lundberg, D.S. et al. Commensal *Pseudomonas* strains facilitate protective response against pathogens in the host plant. Nat Ecol Evol 6, 383–396 (2022). https://doi.org/10.1038/s41559-022-01673-7

38 Wolinska, K. W., Vannier, N., Thiergart, T., Pickel, B., Gremmen, S., Piasecka, A., Piślewska-Bednarek, M., Nakano, R. T., Belkhadir, Y., Bednarek, P., & Hacquard, S. (2021). Tryptophan metabolism and bacterial commensals prevent fungal dysbiosis in *Arabidopsis* roots. Proceedings of the National Academy of Sciences of the United States of America, 118(49), e2111521118. https://doi.org/10.1073/pnas.2111521118

39 Yardeni, T., Tanes, C. E., Bittinger, K., Mattei, L. M., Schaefer, P. M., Singh, L. N., Wu, G. D., Murdock, D. G., & Wallace, D. C. (2019). Host mitochondria influence gut microbiome diversity: A role for ROS. Science signaling, 12(588), eaaw3159. https://doi.org/10.1126/scisignal.aaw3159

40 Miller, B. M., Liou, M. J., Zhang, L. F., Nguyen, H., Litvak, Y., Schorr, E. M., Jang, K. K., Tiffany, C. R., Butler, B. P., & Bäumler, A. J. (2020). Anaerobic Respiration of NOX1-Derived Hydrogen Peroxide Licenses Bacterial Growth at the Colonic Surface. Cell host & microbe, 28(6), 789–797.e5. https://doi.org/10.1016/j.chom.2020.10.009

41 Expert, D., Patrit, O., Shevchik, V. E., Perino, C., Boucher, V., Creze, C., Wenes, E., & Fagard, M. (2018). Dickeya dadantii pectic enzymes necessary for virulence are also responsible for activation of the Arabidopsis thaliana innate immune system. Molecular plant pathology, 19(2), 313–327. https://doi.org/10.1111/mpp.12522

42 Ma, Z., Song, T., Zhu, L., Ye, W., Wang, Y., Shao, Y., Dong, S., Zhang, Z., Dou, D., Zheng, X., Tyler, B. M., & Wang, Y. (2015). A Phytophthora sojae Glycoside Hydrolase 12 Protein Is a Major Virulence Factor during Soybean Infection and Is Recognized as a PAMP. The Plant cell, 27(7), 2057–2072. https://doi.org/10.1105/tpc.15.00390

43 Wang, Y., Xu, Y., Sun, Y., Wang, H., Qi, J., Wan, B., Ye, W., Lin, Y., Shao, Y., Dong, S., Tyler, B. M., & Wang, Y. (2018). Leucine-rich repeat receptor-like gene screen reveals that Nicotiana RXEG1 regulates glycoside hydrolase 12 MAMP detection. Nature communications, 9(1), 594. https://doi.org/10.1038/s41467-018-03010-8

44 Gui, Y. J., Chen, J. Y., Zhang, D. D., Li, N. Y., Li, T. G., Zhang, W. Q., Wang, X. Y., Short, D. P. G., Li, L., Guo, W., Kong, Z. Q., Bao, Y. M., Subbarao, K. V., & Dai, X. F. (2017). Verticillium dahliae manipulates plant immunity by glycoside hydrolase 12 proteins in conjunction with carbohydrate-binding module 1. Environmental microbiology, 19(5), 1914–1932. https://doi.org/10.1111/1462-2920.13695

45 Nobori, T., Wang, Y., Wu, J. et al. Multidimensional gene regulatory landscape of a bacterial pathogen in plants. Nat. Plants 6, 883–896 (2020). https://doi.org/10.1038/s41477-020-0690-7

46 Teixeira, P. J. P., Colaianni, N. R., Fitzpatrick, C. R., & Dangl, J. L. (2019). Beyond pathogens: microbiota interactions with the plant immune system. Current opinion in microbiology, 49, 7–17. https://doi.org/10.1016/j.mib.2019.08.003

47 Vetter, M., Karasov, T. L., & Bergelson, J. (2016). Differentiation between MAMP Triggered Defenses in Arabidopsis thaliana. PLoS genetics, 12(6), e1006068. https://doi.org/10.1371/journal.pgen

48 Furukawa, T., Inagaki, H., Takai, R., Hirai, H., & Che, F. S. (2014). Two distinct EF-Tu epitopes induce immune responses in rice and Arabidopsis. Molecular plant-microbe interactions: MPMI, 27(2), 113–124. https://doi.org/10.1094/MPMI-10-13-0304-R

49 Lacombe, S., Rougon-Cardoso, A., Sherwood, E., Peeters, N., Dahlbeck, D., van Esse, H. P., Smoker, M., Rallapalli, G., Thomma, B. P., Staskawicz, B., Jones, J. D., & Zipfel, C. (2010). Interfamily transfer of a plant pattern-recognition receptor confers broad-spectrum bacterial resistance. Nature biotechnology, 28(4), 365–369. https://doi.org/10.1038/nbt.1613

50 Wang, Y., Garrido-Oter, R., Wu, J. et al. Site-specific cleavage of bacterial MucD by secreted proteases mediates antibacterial resistance in *Arabidopsis*. Nat Commun 10, 2853 (2019). https://doi.org/10.1038/s41467-019-10793-x

51 Wang, W., Yang, J., Zhang, J., Liu, Y. X., Tian, C., Qu, B., Gao, C., Xin, P., Cheng, S., Zhang, W., Miao, P., Li, L., Zhang, X., Chu, J., Zuo, J., Li, J., Bai, Y., Lei, X., & Zhou, J. M. (2020). An Arabidopsis Secondary Metabolite Directly Targets Expression of the Bacterial Type III Secretion System to Inhibit Bacterial Virulence. Cell host & microbe, 27(4), 601–613.e7. https://doi.org/10.1016/j.chom.2020.03.004

52 Lindsey, B. E3rd., Rivero, L., Calhoun, C. S., Grotewold, E., & Brkljacic, J. (2017). Standardized Method for High-throughput Sterilization of Arabidopsis Seeds. Journal of visualized experiments: JoVE, (128), 56587. https://doi.org/10.3791/56587

53 Hinsch M, Staskawicz B. (1996). Identification of a new Arabidopsis disease resistance locus, RPs4, and cloning of the corresponding avirulence gene, avrRps4, from Pseudomonas syringae pv. pisi. Mol Plant Microbe Interact. 9(1):55-61. doi: 10.1094/mpmi-9-0055. PMID: 8589423.

54 Ayumi Matsumoto, Titus Schlüter, Katharina Melkonian, Atsushi Takeda, Hirofumi Nakagami, Akira Mine. (2022). A versatile Tn7 transposon-based bioluminescence tagging tool for quantitative and spatial detection of bacteria in plants, Plant Communications, Volume 3, Issue 1, 2022, 100227, ISSN 2590-3462, https://doi.org/10.1016/j.xplc.2021.100227.

55 Smith, J.M., Heese, A. (2014). Rapid bioassay to measure early reactive oxygen species production in *Arabidopsis* leave tissue in response to living *Pseudomonas syringae*. Plant Methods 10, 6. https://doi.org/10.1186/1746-4811-10-6

56 Merrell, D.S., Hava, D.L. and Camilli, A. (2002), Identification of novel factors involved in colonization and acid tolerance of *Vibrio cholerae*. Molecular Microbiology, 43: 1471–1491. https://doi.org/10.1046/j.1365-2958.2002.02857.x

57 Kvitko, B. H., & Collmer, A. (2011). Construction of Pseudomonas syringae pv. tomato DC3000 mutant and polymutant strains. Methods in Molecular Biology (Clifton,N.J.), 712, 109–128.

58 Gibson, D. G., Young, L., Chuang, R. Y., Venter, J. C., Hutchison, C. A., & Smith, H. O. (2009). Enzymatic assembly of DNA molecules up to several hundred kilobases. Nature Methods, 6(5), 343–345.

59 Kessler, B., V. de Lorenzo, and K. N. Timmis. (1992). A general system to integrate lacZ fusions into the chromosomes of gram-negative eubacteria: regulation of the Pm-promotor of the Tol-plasmid studied with all controlling elements in monocopy. Mol. Gen. Genet. 233:293–301.

60 Wengelnik, K., Marie, C., Russel, M., & Bonas, U. (1996). Expression and localization of HrpA1, a protein of Xanthomonas campestris pv. vesicatoria essential for pathogenicity and induction ofthe hypersensitive reaction. Journal of bacteriology, 178(4), 1061–1069. https://doi.org/10.1128/jb.178.4.1061-1069.1996

61 Nobori, T., & Tsuda, K. (2018). *In planta* Transcriptome Analysis of *Pseudomonas syringae*. Bio-protocol, 8(17), e2987. https://doi.org/10.21769/BioProtoc.2987

62 Liao Y, Smyth GK, Shi W (2019). “The R package Rsubread is easier, faster, cheaper and better for alignment and quantification of RNA sequencing reads.” Nucleic Acids Research, 47, e47. doi: 10.1093/nar/gkz114.

63 Love MI, Huber W, Anders S (2014). “Moderated estimation of fold change and dispersion for RNA-seq data with DESeq2.” Genome Biology, 15, 550. doi: 10.1186/s13059-014-0550-8.

64 Ritchie ME, Phipson B, Wu D, Hu Y, Law CW, Shi W, Smyth GK (2015). “limma powers differential expression analyses for RNA-sequencing and microarray studies.” Nucleic Acids Research, 43(7), e47. doi: 10.1093/nar/gkv007.

65. Storey JD, Bass AJ, Dabney A, Robinson D (2022). qvalue: Q-value estimation for false discovery rate control. R package version 2.30.0, http://github.com/jdstorey/qvalue.

66 R Core Team (2013). R: A language and environment for statistical computing. R Foundation for Statistical Computing, Vienna, Austria. ISBN 3–900051-07-0, http://www.R-project.org/.

66 Charrad, M., Ghazzali, . N., Boiteau, V., & Niknafs, A. (2014). NbClust: An R Package for Determining the Relevant Number of Clusters in a Data Set. Journal of Statistical Software, 61(6), 1–36. https://doi.org/10.18637/jss.v061.i06.

68 Struyf A, Hubert M, Rousseeuw P (1997). “Clustering in an Object-Oriented Environment.” Journal of Statistical Software. doi:10.18637/jss.v001.i04.

69 Gu Z (2022). “Complex Heatmap Visualization.” iMeta. doi: 10.1002/imt2.43

70 Wu T, Hu E, Xu S, Chen M, Guo P, Dai Z, Feng T, Zhou L, Tang W, Zhan L, Fu x, Liu S, Bo X, Yu G (2021). “clusterProfiler 4.0: A universal enrichment tool for interpreting omics data.” The Innovation, 2(3), 100141. doi: 10.1016/j.xinn.2021.100141

71. Mendiburu F and Yaseen M. (2020). agricolae: Statistical Procedures for Agricultural Research. R package version 1.4.0, https://myaseen208.github.io/agricolae/https://myaseen208.github.io/agricolae/project.org/package=agricolae.

72 Drula, E., Garron, M. L., Dogan, S., Lombard, V., Henrissat, B., & Terrapon, N. (2022). The carbohydrate-active enzyme database: functions and literature. Nucleic acids research, 50(D1), D571.D577. https://doi.org/10.1093/nar/gkab1045

73 Cantalapiedra, C. P., Hernandez-Plaza, A., Letunic, I., Bork, P., & Huerta-Cepas, J. (2021). eggNOG-mapper v2: Functional Annotation, Orthology Assignments, and Domain Prediction at the Metagenomic Scale. Molecular biology and evolution, 38(12), 5825.5829. https://doi.org/10.1093/molbev/msab293

